# Identification of two pyruvate transporters in *Salmonella enterica* serovar Typhimurium and their biological relevance

**DOI:** 10.1101/2022.05.13.491600

**Authors:** Stephanie Paulini, Florian D. Fabiani, Anna S. Weiß, Ana Laura Moldoveanu, Sophie Helaine, Bärbel Stecher, Kirsten Jung

**Affiliations:** Department of Microbiology, Ludwig-Maximilians-University Munich, Martinsried, Germany; Bayer AG, Berlin, Germany; Max von Pettenkofer Institute of Hygiene and Medical Microbiology, Faculty of Medicine, Ludwig-Maximilians-University Munich, Munich, Germany; MRC Centre for Molecular Bacteriology and Infection, Imperial College London, London, UK; Department of Microbiology, Harvard Medical School, Boston, United States; German Center for Infection Research (DZIF), partner site LMU Munich, Munich, Germany

**Keywords:** *Salmonella* Typhimurium, pyruvate transporter, chemotaxis, oxidative stress, persistence

## Abstract

Pyruvate (CH_3_COCOOH) is the simplest of the alpha-keto acids and is at the interface of several metabolic pathways both in prokaryotes and eukaryotes. In an amino acid-rich environment, fast-growing bacteria excrete pyruvate instead of completely metabolizing it. The role of pyruvate uptake in pathological conditions is still unclear. In this study, we identified two pyruvate-specific transporters, BtsT and CstA, in *Salmonella enterica* serovar Typhimurium (*S*. Typhimurium). Expression of *btsT* is induced by the histidine kinase/response regulator system BtsS/BtsR upon sensing extracellular pyruvate (threshold 200 μM), whereas expression of *cstA* is maximal in the stationary phase. Both pyruvate transporters were found to be important for the uptake of this compound, but also for chemotaxis to pyruvate, survival under oxidative and nitrosative stress, and persistence of *S*. Typhimurium in response to gentamicin. Compared with the wild-type, the Δ*btsT*Δ*cstA* mutant has disadvantages in antibiotic persistence in macrophages, as well as in colonization and systemic infection in gnotobiotic mice. These data demonstrate the surprising complexity of the two pyruvate uptake systems in *S*. Typhimurium.

## INTRODUCTION

Pyruvate is a primary metabolite of central importance in all living cells. It is the end product of glycolysis and can enter the tricarboxylic acid cycle via acetyl-CoA under aerobic conditions; however, it can also be reduced to lactate under anaerobic conditions. Moreover, it is used as a precursor for the production of amino acids, fatty acids, and sugars. Bacteria tightly control intracellular pyruvate levels, which are normally approximately 40 μM. In an amino acid-rich environment, fast-growing bacteria excrete pyruvate instead of metabolizing it completely, a phenomenon known as overflow metabolism, and take it up again later (Behr et al., 2017a; Chubukov et al., 2014; Paczia et al., 2012; Yasid et al., 2016).

Pyruvate also scavenges reactive oxygen species (ROS). It inactivates hydrogen peroxide (H_2_O_2_) by being oxidized and rapidly decarboxylated (Constantopoulos & Barranger, 1984; Kładna et al., 2015; Varma et al., 2003). Therefore, the secretion of pyruvate can also be seen as an antioxidant defense mechanism (O’Donnell-Tormey et al., 1987). The role of pyruvate in the inactivation of ROS is important for the resuscitation of viable but non-culturable (VBNC) bacteria. Pyruvate is required to “wake up” cells from this dormant state and re-enter culturability (Dong et al., 2020; Göing et al., 2021; Mizunoe et al., 1999; Vilhena et al., 2019). *S*. Typhimurium was effectively resuscitated from the VBNC state using pyruvate (Liao et al., 2018).

Several reports have demonstrated the importance of pyruvate as focal point in metabolism and in virulence control of pathogens, such as *Yersinia pseudotuberculosis, S*. Typhimurium, *Listeria monocytogenes*, and *Vibrio parahaemolyticus* (Abernathy et al., 2013; Bücker et al., 2014; Schär et al., 2010; van Doorn et al., 2021; Xie et al., 2019). *Pseudomonas aeruginosa, Staphylococcus aureus*, and *Chlostridium difficile* require extracellular pyruvate for biofilm formation (Goodwine et al., 2019; Petrova et al., 2012; Tremblay et al., 2021). Mammalian apoptotic cells also release pyruvate, which has been shown to promote the growth of *S*. Typhimurium (Anderson et al., 2021). This suggests an important role for pyruvate in host inflammation and infection.

In *E. coli*, BtsT and CstA have been characterized as substrate-specific pyruvate transporters, and a deletion mutant of these two transporter genes and the gene *yhjX* has lost the ability to grow on pyruvate, indicating that YhjX might also be a pyruvate transporter (Gasperotti et al., 2020; Hwang et al., 2018; Kristoficova et al., 2018). *btsT* and *yhjX* are activated by the histidine kinase/response regulator systems BtsS/BtsR and YpdA/YpdB (PyrS/PyrR), respectively, when the cells sense pyruvate (Behr et al., 2014; Behr et al., 2017b; Fried et al., 2013; Hwang et al., 2018), whereas *cstA* is induced by nutrient limitation in the stationary phase (Gasperotti et al., 2020; Schultz & Matin, 1991). There are some monocarboxylate transporters that have broader substrate specificity and can also transport pyruvate: MctP in *Rhizobium leguminosarum* (Hosie et al., 2002), MctC in *Corynebacterium glutamicum* (Jolkver et al., 2009), PftAB in *Bacillus subtilis*, which is activated by the LytS/LytT two-component system (Charbonnier et al., 2017), and LrgAB in *Streptococcus mutans* (Ahn et al., 2019).

The enteric pathogen *Salmonella* is one of the leading causes of acute diarrheal disease, which affects more than 2 billion people worldwide each year (Popa & Papa, 2021). *S*. Typhimurium was shown to excrete and likewise to reclaim pyruvate (Behr et al., 2017a), as well as to grow on pyruvate as the sole carbon source (Christopherson et al., 2008), but no pyruvate transporter has been characterized yet. Homologs of *E. coli* genes *btsT* and *cstA* are found in *S*. Typhimurium, which we designate as *btsT* (locus tag SL1344_4463), previously known as *cstA1* or *yjiY*, and *cstA* (locus tag SL1344_0588). Both genes have been previously described to be involved in peptide utilization and in the colonization of *C. elegans* and mice (Garai et al., 2016; Tenor et al., 2004). Wong et al. (2013) investigated the histidine kinase/response regulator system BtsS/BtsR (previously known as YehU/YehT) in *S*. Typhi and Typhimurium and identified *btsT* as a predominantly regulated gene. Finally, an unusually high number of mutations over lineage development accumulated in the *btsSR* operon (Holt et al., 2008), suggesting that this system is targeted by adaptive evolution and is therefore of potential significance for the pathogen.

Here, we characterized BtsT and CstA as pyruvate transporters of *S*. Typhimurium and evaluated their importance for the pathogen *in vitro* and *in vivo.*

## RESULTS AND DISCUSSION

### S. *Typhimurium possesses two pyruvate transporters, BtsT and CstA*

Based on homology search, *S.* Typhimurium has two genes coding for putative pyruvate transporters: *btsT* (locus tag SL1344_4463) codes for a 77 kDa transporter protein that shares 96.6% identity with the *E. coli* BtsT, according to the online tool Clustal Omega (Sievers et al., 2011). *S.* Typhimurium *cstA* (locus tag SL1344_0588) codes for a 75 kDa transporter protein that shares 97.1% identity with the *E. coli* CstA. The genetic contexts of *btsT* and *cstA* are illustrated in Figure 1A. Both transporters belong to the CstA family (transporter classification: [TC] 2. A.114) (Saier et al., 2021) with at least 16 predicted transmembrane domains (UniProt, 2021) and share 60.5 % identity and 24.1% similarity with each other at 97.2% coverage, as illustrated in Figure 1B.

**Figure 1.**
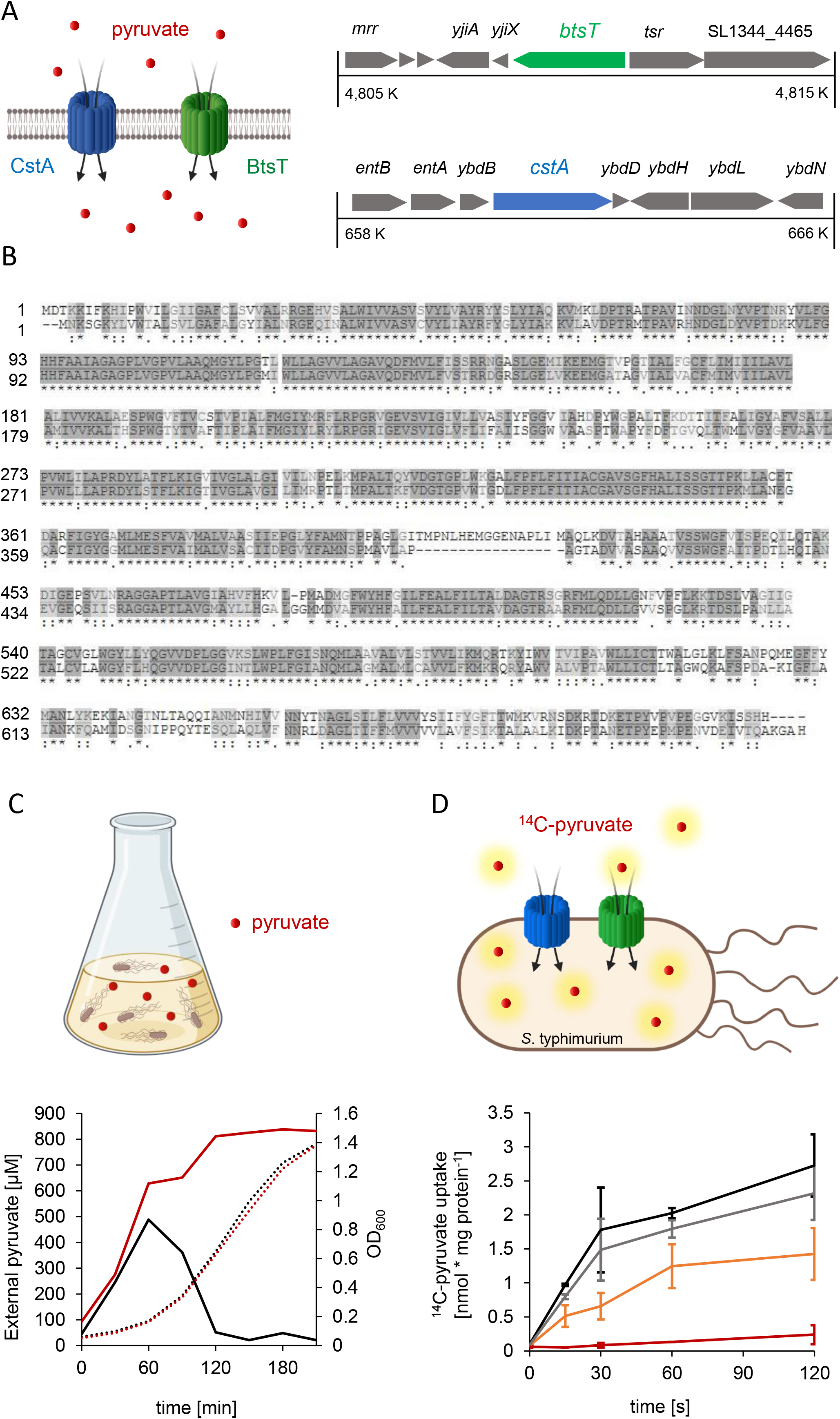
*S.* Typhimurium possesses two pyruvate transporters, BtsT and CstA. **A)** Schematic illustration of the two transporters BtsT and CstA in *S*. Typhimurium responsible for the uptake of pyruvate and the genetic context of their genes [*btsT* (SL1344_4463), *cstA* (SL1344_0588)]. **B)** Protein sequence alignment of BtsT (upper line) and CstA (lower line), created with the online tool Clustal Omega. **C)** Alterations of the pyruvate concentration in LB medium (solid lines) owing to overflow and uptake during growth (dotted lines) of *S*. Typhimurium SL1344 wild-type (black) and Δ*btsT*Δ*cstA* mutant (red). Samples were taken every 20 min. **D)** Time course of [^14^C]-pyruvate (10 μM) uptake by intact cells at 18°C: SL1344 wild-type (black) and Δ*btsT*Δ*cstA* mutant (red). Error bars represent the standard deviations of the mean of three individual experiments. All illustrations were created with BioRender.

Gamma-proteobacteria excrete pyruvate when grown in amino acid-rich media, such as LB, owing to an overflow metabolism (Behr et al., 2017a; Chubukov et al., 2014; Paczia et al., 2012). We measured the external pyruvate concentration during the growth of wild-type *S.* Typhimurium in LB (Figure 1C). At the beginning of exponential growth, the pyruvate concentration in the LB medium increased from 60 μM to 490 μM, followed by a rapid decrease back to the initial pyruvate concentration. For the double deletion mutant Δ*btsT*Δ*cstA*, we monitored the same pyruvate excretion as the wild-type but did not observe any subsequent decrease in the external pyruvate concentration; on the contrary, the concentration increased further, reaching a plateau at 800 μM (Figure 1C). This indicates that the Δ*btsT*Δ*cstA* mutant did not reclaim pyruvate after excretion, which then accumulated in the medium.

To further confirm that BtsT and CstA are the only pyruvate transporters in *S.* Typhimurium, we performed transport experiments with radiolabeled pyruvate and intact cells. To avoid rapid metabolization, all assays were performed at 18°C. For wild-type *S.* Typhimurium, we monitored the uptake of radiolabeled pyruvate over time (Figure 1D), with an initial uptake rate of 3.5 nmol per mg protein per minute and a maximal uptake of 2.7 nmol per mg protein, whereas for the double deletion mutant Δ*btsT*Δ*cstA* no transport of radiolabeled pyruvate was observed (Figure 1D). Both single deletion mutants were able to take up pyruvate, but at a decreased rate (Figure 1D). This shows that BtsT and CstA transport pyruvate into *S.* Typhimurium cells.

### *Expression of* btsT *is activated by the histidine kinase response regulator system BtsS/BtsR in the presence of pyruvate, whereas expression of* cstA *is dependent on the growth phase*

To investigate growth-dependent *btsT* and *cstA* activation, we used luciferase-based reporter strains (Figure 2A). Cells were grown in LB medium and a sharp *btsT* expression peak was observed at the beginning of the exponential growth phase (Figure 2C). This expression pattern is very similar to that observed in *E. coli* (Behr et al., 2014; Behr et al., 2017b). Expression of *cstA* started at the beginning of the stationary phase (Figure 2D). This expression pattern was similar for *E. coli cstA* and is explained by the induction of *cstA* under nutrient limitation as an effect of at least two regulators: cAMP-CRP and Fis (Gasperotti et al., 2020; Schultz & Matin, 1991).

**Figure 2.**
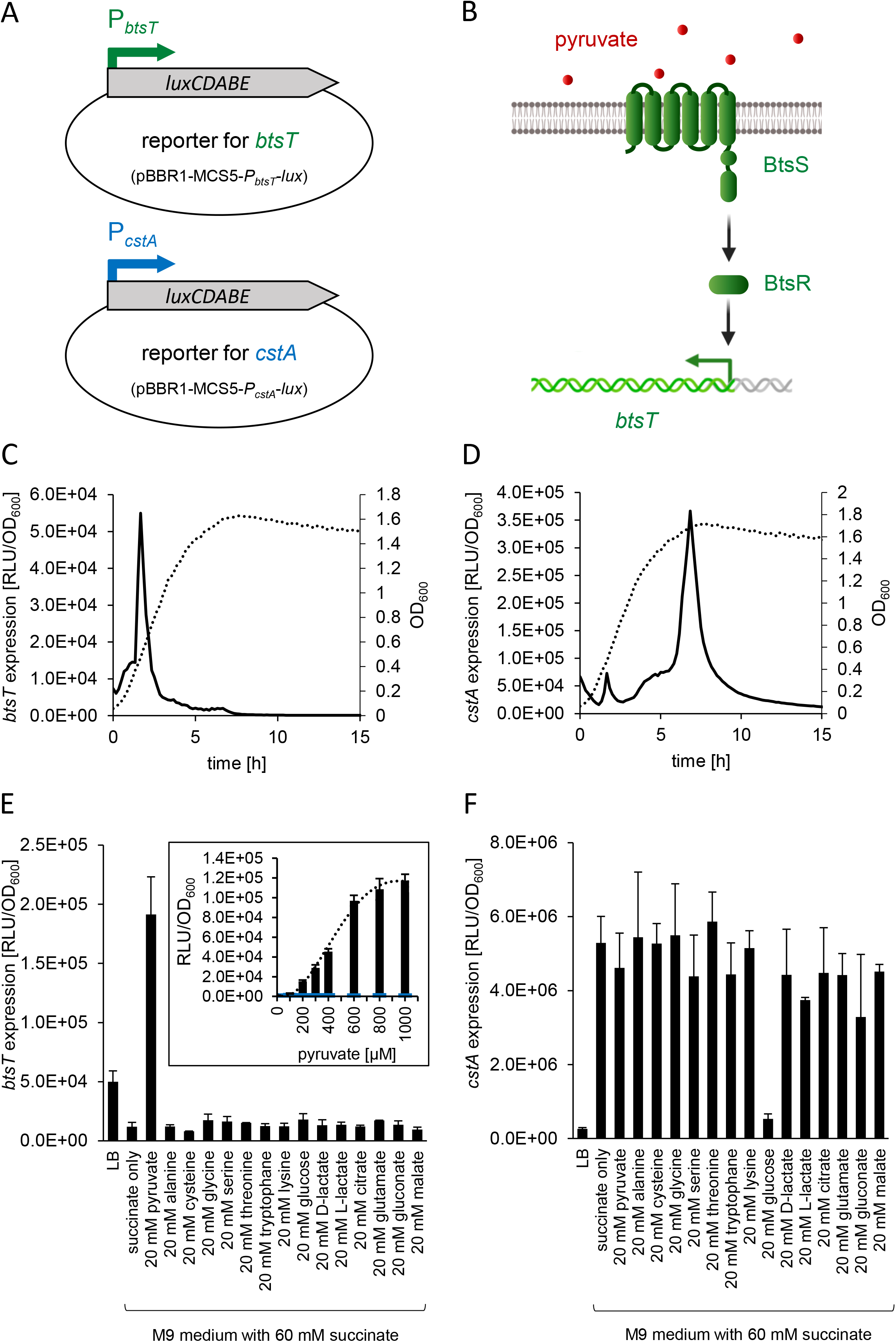
Expression of *btsT* and *cstA* in *S.* Typhimurium. **A)** Schematic illustration of the luciferase-based, low copy reporter plasmids to monitor *btsT* (pBBR1-MCS5-*PbtsT*-*lux*) and *cstA* (pBBR1-MCS5-*PcstA*-*lux*) expression. **B)** Schematic illustration of the two-component system BtsS/BtsR in *S*. Typhimurium, with the histidine kinase BtsS sensing pyruvate and the response regulator BtsR inducing *btsT*. **C)** Expression of *btsT* in *S*. Typhimurium SL1344 (pBBR1-MCS5-*P_btsT_*-*lux*) during growth in LB medium at 37°C. Luminescence (RLU normalized to OD_600_ = 1) (solid line) and growth (OD_600_) (dotted line) were measured over time in a plate reader. The graphs show the means of three independent replicates; the standard deviations were below 10%. **D)** Expression of *cstA* in S. Typhimurium SL1344 (pBBR1-MCS5-*PcstA*-*lux*) during growth in LB medium. Experimental set-up as in **C**; OD_600_ (dotted line), RLU per OD_600_ (solid line). The graphs show the means of three independent replicates; the standard deviations were below 10%. **E)** Expression of *btsT* in SL1344 (pBBR1-MCS5-*PbtsT*-*lux*) grown in M9 minimal medium supplemented with 60 mM succinate and the indicated C-sources, each at 20 mM. Experimental set-up as in **C.** The maximal RLUs per OD_600_ served as the measure for *btsT* expression. The value of the basal activation in the presence of succinate was subtracted. Inset: Expression of *btsT* in wild-type (black) or Δ*btsSR* (blue) cells as a function of pyruvate concentration. Cells were grown in M9 minimal medium with 60 mM succinate and different concentrations of pyruvate. The value of the basal activation in the presence of succinate was subtracted. **F)** Expression of *cstA* in SL1344 (pBBR1-MCS5-*PcstA-lux*) grown in M9 minimal medium. Experimental set-up as in **E**. **E**, **F**, Error bars represent the standard deviations of the mean of three independent replicates. Illustrations were partly created with BioRender.

We then measured the expression of *btsT* and *cstA* in cells grown in minimal medium containing different carbon (C) sources. Expression of *btsT* was exclusively activated in cells grown in minimal medium with pyruvate as the sole C-source and barely in the presence of other compounds, such as amino acids or different carboxylic acids (Figure 2E).

In *E. coli*, *btsT* expression was shown to be activated by the LytS/LytTR-type two-component system BtsS/BtsR upon sensing pyruvate (Behr et al., 2017b; Kraxenberger et al., 2012). Wong et al. (2013) also showed the activation of *btsT* by BtsS/BtsR in *S*. Typhimurium, but did not identify the responsible stimulus. We created deletion mutants of the homologous genes of this two-component system in *S.* Typhimurium SL1344 (gene *btsS*, locus tag SL1344_2137, coding for the histidine kinase BtsS, and gene *btsR*, locus tag SL1344_2136, coding for the response regulator BtsR) to test whether the system is required for *btsT* activation. Indeed, we could not detect *btsT* expression in cells lacking *btsSR* (inset panel in Figure 2E).

To further analyze the activation of *btsT* expression by pyruvate, cells were grown in minimal medium with different pyruvate concentrations (and 60 mM succinate as the basic C-source for growth, for which the activation value was subtracted). We monitored the concentration-dependent activation of *btsT* by pyruvate, with a threshold concentration of 200 μM required for induction and saturation of *btsT* expression at approximately 1 mM (inset panel in Figure 2E). The pyruvate concentration that resulted in half-maximal *btsT* expression was estimated to be 450 μM. We conclude that the transcriptional activation of *btsT* in *S.* Typhimurium follows the same pattern as that in *E. coli*, and that pyruvate sensing BtsS/BtsR activates *btsT* expression to mediate rapid uptake of the compound by BtsT (illustrated in Figure 2B).

In contrast, high expression of *cstA* was observed in cells independent of the C-source (Figure 2F). Expression of *cstA* was lower when cells were grown in amino acid-rich LB medium and in glucose-containing minimal medium, indicating a control by nutrient availability and catabolite repression. Indeed, *cstA* expression was always highest in stationary phase cells.

We also analyzed the expression of *btsT* and *cstA* in mutants with deletions of either *btsT, cstA* or both *btsT* and *cstA* grown in LB medium (Figure S1). We found a 12-fold upregulation of *btsT* in the Δ*btsT* mutant and a 50-fold upregulation in the Δ*btsT*Δ*cstA* mutant compared to that in the wild-type (Figure S1C). A similar feedback regulation was observed for *btsT* in the pathogen *Vibrio campbellii* (Göing et al., 2021), but the exact mechanism is unknown. In Δ*cstA* cells, the expression pattern of *btsT* over time was the same as that in the wild-type (Figure S1C). Moreover, the pattern and level of *cstA* expression in all mutants were identical to those in the wild-type (Figure S1B).

We conclude that *btsT* expression is activated by pyruvate, whereas *cstA* expression is induced in stationary phase and repressed by glucose.

### Pyruvate uptake by BtsT and CstA is required for growth on pyruvate and chemotaxis to pyruvate

In the next step, we investigated the biological impact of pyruvate uptake by BtsT and CstA in *S.* Typhimurium through phenotypical characterization of the Δ*btsT*Δ*cstA* mutant in comparison to the wild-type.

The *S.* Typhimurium Δ*btsT*Δ*cstA* deletion mutant was unable to grow on pyruvate as the sole C-source (Figure 3) but grew on other C-sources such as glucose or in complex media such as LB (Figure S2A). Full complementation of the double deletion mutant Δ*btsT*Δ*cstA* was achieved by expressing both *btsT* and *cstA in trans* (Figure 3). The single deletion mutants Δ*bstT* and Δ*cstA* were able to grow on pyruvate, albeit not as well as the wild-type (Figure S2B). Expression of *btsT* alone was sufficient to restore growth to almost the wild-type level, whereas expression of *cstA* alone could only partially restore growth (Figure 3).

**Figure 3.**
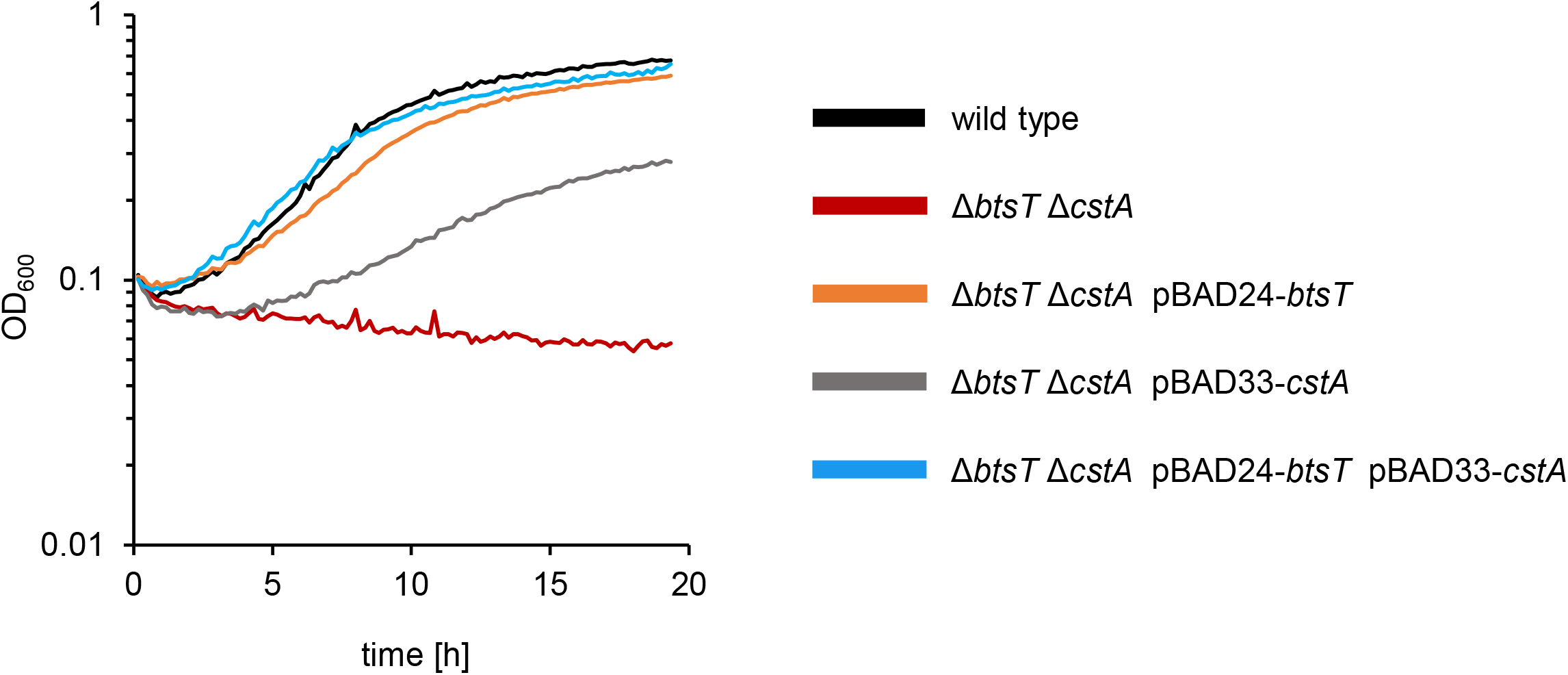
*S*. Typhimurium mutant Δ*btsT*Δ*cstA* is unable to grow on pyruvate. SL1344 wild-type and Δ*btsT*Δ*cstA* mutant harboring the indicated plasmid(s) were grown in M9 minimal medium with 60 mM pyruvate in a plate reader at 37°C.

We then analyzed the chemotactic behavior of wild-type and Δ*btsT*Δ*cstA S.* Typhimurium cells using the plug-in-pond assay (Darias et al., 2014), in which cells are mixed with soft agar and poured into a petri dish containing agar plugs with potential attractants (Figure 4A). When the cells respond chemotactically to an attractant, a ring of clustered cells is visible around the agar plug. For wild-type *S.* Typhimurium, we observed chemotaxis to pyruvate by a clearly visible ring of accumulating cells (Figure 4A). In contrast, no ring was found in the Δ*btsT*Δ*cstA* mutant, indicating the loss of chemotaxis to pyruvate. This phenotype could be complemented by expressing *btsT* and *cstA in trans* (Figure S3).

**Figure 4.**
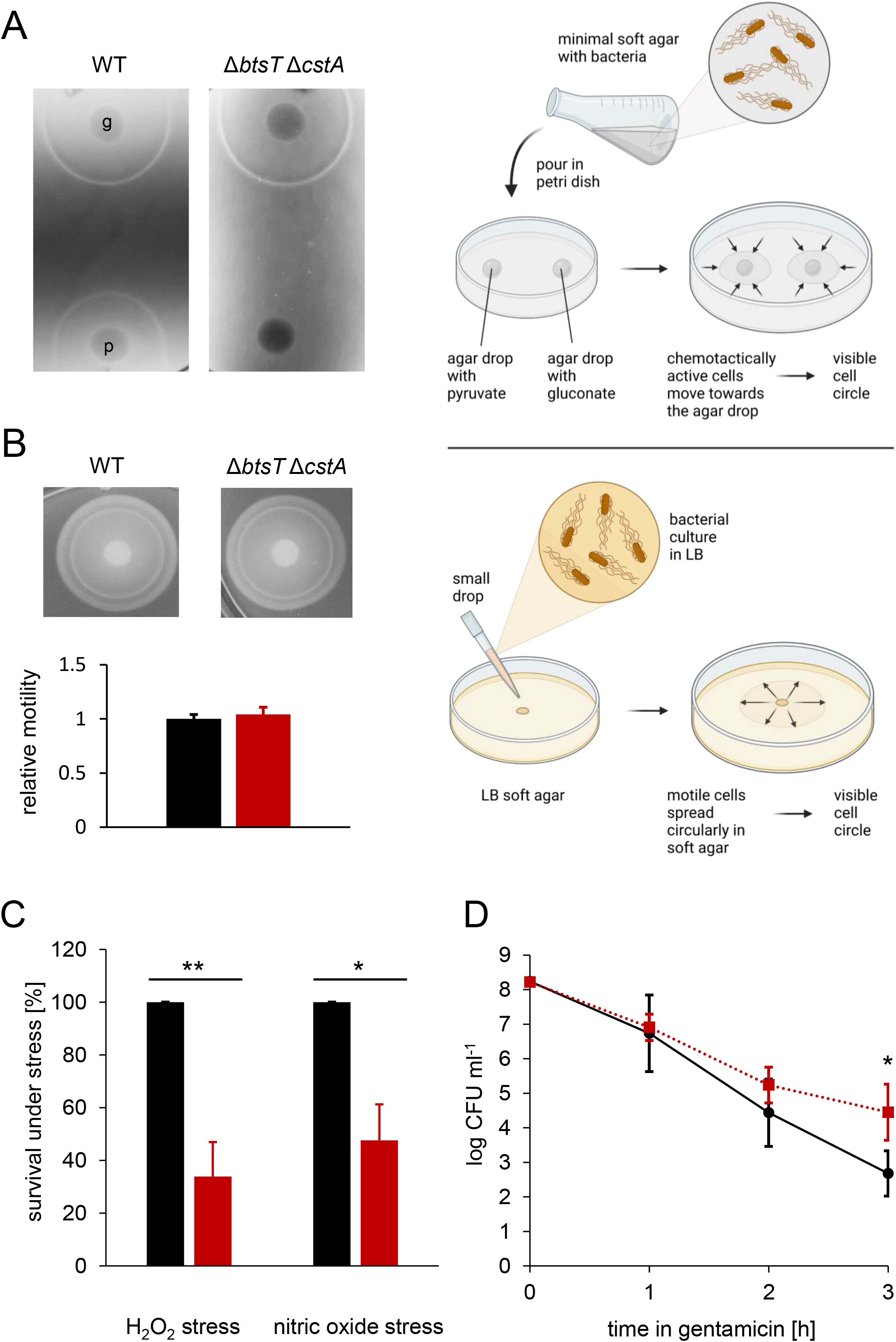
*In vitro* phenotypes of *S.* Typhimurium Δ*btsT*Δ*cstA* mutant. **A)** Chemotaxis assay with schematic illustration: Chemotaxis was tested by mixing SL1344 wild-type (left) and Δ*btsT*Δ*cstA* (right) cells with 0.3% (wt/vol) M9 soft agar and pouring them over 1.5% (wt/vol) M9 agar plugs containing either 50 mM gluconate (g) or 50 mM pyruvate (p). Plates were incubated at 37°C for 4 h, and the pictures are representative of three independent experiments. **B)** Swimming motility assay with schematic illustration: Motility of SL1344 wild-type (left, black) and Δ*btsT*Δ*cstA* (right, red) cells was tested by spotting equal numbers of cells on 0.3% (wt/vol) LB soft agar, incubating the plates at 37°C for 3 h and measuring the cell ring diameter with the software ImageJ. Images of rings are representative of four independent experiments and relative motility was determined in relation to the mean diameter of the wild-type ring. **C)** Oxidative and nitrosative stress tests: SL1344 wild-type (black) and Δ*btsT*Δ*cstA* (red) cells were grown in LB medium to OD_600_ = 1.2, split in two groups and exposed to 12.5 mM H_2_O_2_ or 250 μM spermine NONOate or H_2_O as control. After 20 min of incubation, catalase was added to the H_2_O_2_ treated group and cells were plated in dilutions on LB plates to determine CFU. Survival under stress was calculated as the percentage of CFU in relation to the control condition, and wild-type values were set to 100%. Error bars represent the standard deviations of the mean of three independent experiments. **D)** Formation of antibiotic-induced persister cells: SL1344 wild-type (black, circles) and Δ*btsT*Δ*cstA* (red, squares) cells were grown in LB medium to OD_600_ = 1.2 and diluted to OD_600_ = 0.05 into fresh LB containing 50 μg/ml gentamicin. Every hour, cells were plated in dilutions on LB plates to determine CFU. Error bars represent the standard deviations of the mean of three independent experiments. \**p* < 0.05, \*\**p* < 0.01. All illustrations were created with BioRender.

To ensure that this defect did not result from impaired swimming motility, we analyzed the swimming motility of wild-type and mutant cells in LB soft agar, as illustrated in Figure 4B, and could not see any difference; both strains moved in the soft agar circularly away from the inoculation spot, where the cells had been dropped before, and formed visible rings of cells after 3 h of incubation (Figure 4B). The measurement of the ring sizes clearly shows that both strains could swim to the same extent. This finding is important, as it was previously claimed that motility and flagella biosynthesis are impaired in an *S*. Typhimurium Δ*btsT* mutant (Garai et al., 2016). We could not confirm these previously published results, neither for the double deletion mutant Δ*btsT*Δ*cstA* (Figure 4B) nor for the single deletion mutants Δ*btsT* or Δ*cstA* (Figure S4). It should be noted that Garai et al. (2016) did not report the successful complementation of deletion mutants.

Importantly, chemotaxis to other substances, such as gluconate, was not affected by deletions of *btsT* and *cstA*, as the double deletion mutant showed the same ring of accumulated cells as the wild-type (Figure 4A). These results indicate that pyruvate uptake is necessary for chemotaxis to pyruvate, leading to the conclusion that the chemotactic response must be activated by intracellular pyruvate. Similarly, for other gamma-proteobacteria it was described previously that the deletion of pyruvate transporter gene(s) impairs chemotaxis to pyruvate (Gasperotti et al., 2020; Göing et al., 2021). In *E. coli*, it has been shown that the phosphotransferase system (PTS) can sense pyruvate inside cells and that signals from the PTS are transmitted linearly to the chemotaxis system (Neumann et al., 2012; Somavanshi et al., 2016). Thus, we conclude that *S.* typhimurium must take up pyruvate, and the PTS monitors intracellular pyruvate levels via the ratio of pyruvate to phosphoenolpyruvate to trigger a chemotactic response to this compound.

### Pyruvate uptake is important to survive oxidative stress, nitrosative stress, and antibiotic treatment

The production of ROS and nitric oxide (NO) is an important defense mechanism of the host to control the proliferation of intracellular pathogens, such as *S*. Typhimurium (Richardson et al., 2011). Pyruvate is a known scavenger of ROS (Constantopoulos & Barranger, 1984; Kładna et al., 2015; Varma et al., 2003). Thus, we analyzed the importance of pyruvate uptake by *S.* Typhimurium under ROS and NO stress. We challenged wild-type and Δ*btsT*Δ*cstA S.* Typhimurium by exposing cells to hydrogen peroxide (H_2_O_2_) and nitrosative stress (NO) for 20 min. We found that the double mutant had a clear disadvantage compared to the wild-type (Figure 4C). Only half as many Δ*btsT*Δ*cstA* as wild-type cells were able to survive these stressful conditions, indicating that pyruvate uptake is important for *S.* Typhimurium to cope with oxygen and nitric radicals. We conclude that intracellular pyruvate is required as a ROS scavenger and to compensate for the metabolic defects caused by NO. Kröger et al. (2013) found a slight upregulation of *btsT* and *cstA* under oxidative stress.

We also compared *S.* Typhimurium double pyruvate transporter mutant with the wild-type under antibiotic stress. Bacterial persisters survive exposure to antibiotics in laboratory media owing to their low metabolic activity and low growth rate (Balaban et al., 2004; Lewis, 2010). We exposed wild-type and Δ*btsT*Δ*cstA S.* Typhimurium cells to gentamicin (50 μg/mL) and monitored the number of colony forming units (CFU) over time. Only cells able to survive this stress form CFU. We observed a steep initial decrease in CFU for both strains, followed by a slower killing rate in the case of the mutant, which typically reveals the persister fraction of the population (Figure 4D). We hypothesize that the deficit in pyruvate uptake results in cells with lower metabolic activity, which are less harmed by antibiotic stress. Similarly, in *E. coli*, a pyruvate sensing network that tightly regulates the expression of two pyruvate transporters is important for balancing the physiological state of the entire population and increasing the fitness of single cells (Vilhena et al., 2018). An *E. coli* mutant that is unable to produce the two major pyruvate transporters forms more persister cells than the wild-type (Vilhena et al., 2018). We also quantified the persister fractions surviving other antibiotics, such as ampicillin and cefotaxime, in *S.* Typhimurium, but did not find any difference between the wild-type and mutant (data not shown).

### Pyruvate uptake is important to recover from intra-macrophage antibiotic treatment

The facultative intracellular pathogen *S.* Typhimurium forms non-growing antibiotic persisters at high levels within macrophages (Helaine et al., 2014), which have a different physiological state than persisters formed *in vitro* (Stapels et al., 2018). Therefore, we investigated whether pyruvate uptake plays a role in intra-macrophage antibiotic survival. As illustrated in Figure 5A, macrophages were infected with wild-type or Δ*btsT*Δ*cstA S.* Typhimurium cells, and after 30 min of incubation, the bacteria were recovered following lysis of half of the infected macrophages, and the number of surviving bacteria was determined by plating and counting CFU. The other half of the infected macrophages was challenged with cefotaxime for 24 h. After this treatment, the number of bacteria was determined, as described above. By comparing the number of CFU before and after cefotaxime treatment, the survival of *S.* Typhimurium cells in the macrophages during antibiotic stress was calculated.

**Figure 5.**
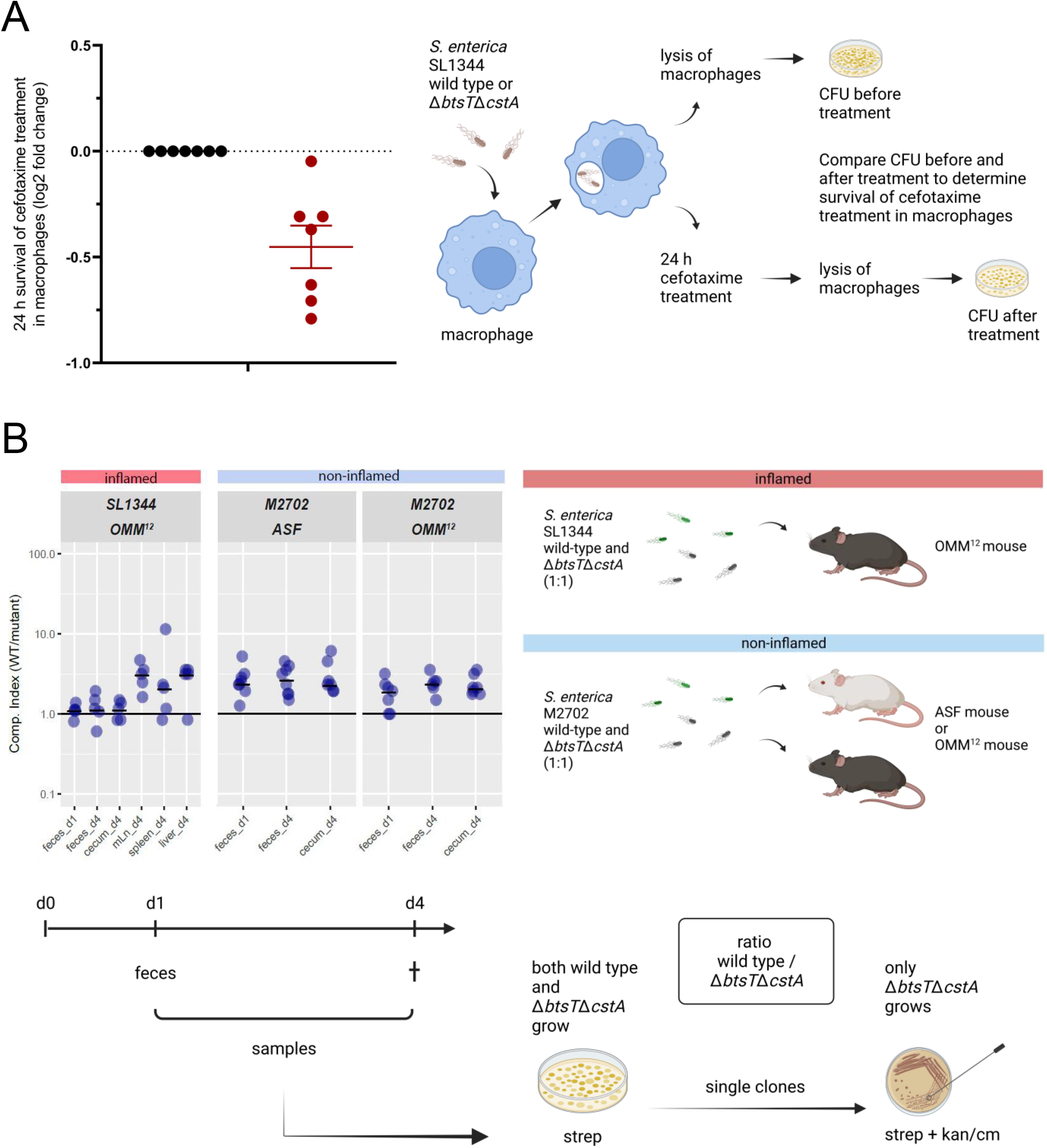
*In vivo* phenotypes of *S.* Typhimurium Δ*btsT*Δ*cstA* mutant. **A)** Intra-macrophage antibiotic survival assay with schematic illustration: Bone marrow derived macrophages were infected with either SL1344 wild-type or Δ*btsT*Δ*cstA* stationary phase bacteria. After 30 min, one part of the macrophages was lysed and the recovered bacteria were plated to determine CFUs. The other part of infected macrophages was treated with cefotaxime and incubated for 24 h, followed by macrophage lysis and plating to determine CFUs. The number of CFU after antibiotic treatment was set in relation to the number of CFU prior to antibiotic treatment. The 24 h antibiotic survival was then expressed as a fold-change of wild-type values. **B)** Competition assay in gnotobiotic mice with schematic illustration: OMM^12^ or ASF mice were inoculated with both wild-type and Δ*btsT*Δ*cstA* cells (ratio 1:1) of the virulent strain SL1344 or the avirulent strain M2702, respectively. One day after infection, fecal samples were collected and plated on MacConkey agar with streptomycin, which selects for all *S. enterica* cells owing to natural resistance. Four days after infection, all mice were sacrificed and samples from feces, cecum, lymph nodes, spleen and liver were plated on MacConkey agar with streptomycin. Single colonies were streaked on MacConkey agar with streptomycin plus kanamycin (for SL1344) or chloramphenicol (for M2702) to select for Δ*btsT*Δ*cstA* cells. By this means, the competitive index could be determined, i.e. the ratio between wild-type and Δ*btsT*Δ*cstA* cells. All illustrations were created with BioRender.

The Δ*btsT*Δ*cstA* mutant had impaired survival to cefotaxime treatment within the macrophages compared to the wild-type (Figure 5A). These results show that pyruvate uptake plays a role in *S.* Typhimurium survival in cefotaxime-treated macrophages. The difference between wild-type and mutant cells in the intramacrophage survival assay was rather small. Although the macrophage environment and the *in vitro* conditions are not really comparable, we also measured only a low and homogeneous activation of *btsT* during growth of *S*. Typhimurium in InSPI2 medium (Löber et al., 2006) (Figure S5). For VBNC *E. coli* cells, we have previously shown that pyruvate is the first substrate taken up when cells return to the culturable state (Vilhena et al., 2019), and pyruvate is likewise important for the resuscitation of *S*. Typhimurium (Liao et al., 2018). We propose that the uptake of pyruvate is important for the regrowth of *S*. Typhimurium from the persister state out of macrophages.

### Mutants lacking pyruvate transporters show a slight disadvantage in colonization and systemic infection of gnotobiotic mice

*S*. Typhimurium colonizes the gut of its host, leading to inflammation, but it can also disseminate inside macrophages to other organs and cause systemic infection. In mice infected with *S*. Typhimurium, pyruvate concentrations were found to be significantly higher than those in uninfected mice (Anderson et al., 2021). Therefore, we investigated how the double deletion of pyruvate transporter genes in *S*. Typhimurium affects the colonization of gnotobiotic mice, as illustrated in Figure 5B. First, we used OMM^12^ mice, which stably carry a minimal consortium of 12 bacterial strains (Brugiroux et al., 2016). To reduce colonization resistance and allow infection by *S.* Typhimurium SL1344, OMM^12^ mice were pretreated with streptomycin. In a competition assay, OMM^12^ mice were infected with a 1:1 mixture of both SL1344 wild-type and Δ*btsT*Δ*cstA* mutant. One day after infection, fecal samples were taken, and four days after infection, mice were sacrificed, and samples from the cecum, feces, and different organs were collected. Notably, mice developed gut inflammation owing to infection with virulent *S.* Typhimurium SL1344. To determine the number of *S.* Typhimurium bacteria, samples were plated on streptomycin, an antibiotic which *S.* Typhimurium is resistant to. From these plates, several single clones were picked and streaked on kanamycin to determine the proportion of these cells as Δ*btsT*Δ*cstA* mutants, as only mutant cells carry the kanamycin resistance cassette. Thus, the competitive index, that is the ratio between the wild-type and mutant cells, was determined.

We found that, in all samples, the average competitive index was higher than 1, indicating that more SL1344 wild-type than Δ*btsT*Δ*cstA* cells were present (Figure 5B). In fecal samples, both one day and four days post infection, as well as in cecum samples, the competitive index was just slightly higher than 1, indicating that both the wild-type and mutant colonized equally well. However, in the lymph nodes, spleen, and liver, organs to which *S.* Typhimurium disseminates to cause systemic infection, an average competitive index of approximately 3 indicated a three times higher number of wild-type than mutant cells. These findings indicate that *S*. Typhimurium SL1344 Δ*btsT*Δ*cstA* cells, which cannot take up pyruvate, have a disadvantage in the systemic infection of OMM^12^ mice.

It has been shown that, in mice colonized with a different minimal bacterial consortium, the so-called altered Schaedler flora (ASF mice), more nutrients are available, *S*. Typhimurium *btsT* is upregulated, and no colonization resistance against the pathogen is provided (Eberl et al., 2021). We also investigated the competition between Δ*btsT*Δ*cstA* mutant and wild-type cells in these mice. Infection with SL1344 bacteria induces severe colitis in ASF mice that lack a sufficiently protective microbiota. Therefore, we generated deletions of both *btsT* and *cstA* in a non-virulent *S.* Typhimurium strain, M2702 (lacking the two virulence factors *invG* and *ssaV*), with the final Δ*btsT*Δ*cstA* mutant carrying a chloramphenicol resistance cassette to distinguish it from the wild-type. Competition experiments were performed as previously described and are illustrated in Figure 5B, with three differences compared to infection experiments with the virulent *S.* Typhimurium strain: no antibiotic treatment was carried out before infection and chloramphenicol instead of kanamycin was used to select for the mutant cells. Moreover, no organ samples from the lymph nodes, spleen, or liver were taken, as the non-virulent *S.* Typhimurium M2702 bacteria are able to colonize but not to systemically infect the mice.

In ASF mice, the average competitive index was higher than 1 for all samples (Figure 5B). Approximately three times more wild-type than Δ*btsT*Δ*cstA* mutant cells were counted in fecal and cecum samples. This indicates that avirulent *S.* Typhimurium bacteria unable to take up pyruvate had a disadvantage in colonizing the non-inflamed gut of ASF mice. Although the competitive index numbers were rather subtle, there was a clear difference between the wild-type and the mutant in samples taken from the cecum and feces. This trend was equally observable in the non-inflamed environment of OMM^12^ mice, indicating that the colonization differences of *S*. Typhimurium were not microbiota-dependent.

We assume that for the non-virulent M2702 bacteria, the advantage of the wild-type cells might have already come to light in the gut, as they compete only there with the mutants. For the virulent SL1344 bacteria, in contrast, the advantage of wild-type cells could have led to more cells entering macrophages and traveling to organs such as lymph nodes or the liver. This could explain why the differences between wild-type and mutant cells regarding gut colonization were only found under non-virulent conditions. The microbiota did not show any influence on the competition between wild-type and mutant cells. Another explanation could be that the difference between the avirulent wild-type and mutant resulted from the different environment in the non-inflamed gut, where other nutrients are available. We conclude that pyruvate uptake delivers a small advantage for *S*. Typhimurium in both colonization and – if the cells are able to – systemic infection of gnotobiotic mice.

We expected to see a stronger disadvantage of the *S*. Typhimurium pyruvate transporter mutant in the *in vivo* experiments. However, extreme phenotypes cannot be expected *in vivo* by preventing the uptake of one compound. In macrophages, pyruvate uptake might help deal with oxidative stress, but there are other factors that are important and overlay this effect. In the gut, pyruvate is present, even more in the inflamed gut and during *Salmonella* infection, but the question is, if it is even available and necessary in this state for *S*. Typhimurium, so that it can depict an advantage. The intestine and its microbiome are a complex ecosystem with interaction networks of numerous bacterial communities and metabolites in distinct niches (Gilbert et al., 2018). The minimal bacterial consortia used in this study are still what their name says, minimal, providing at most a model intestinal ecosystem (Clavel et al., 2016), and their metabolic interactions are not yet fully solved (Weiss et al., 2021). We must consider that the importance of pyruvate putatively did not entirely come to light here, and both wild-type and mutant bacteria were not under pressure to give pyruvate uptake a strong impact on fitness and virulence, as they may have been in a more complex community.

## CONCLUSION

This study is the first to describe pyruvate transport in *Salmonella* and its importance for the cells also beyond metabolism. Especially for relevant pathogens, it is very important to gain more and detailed knowledge about how they use specific compounds and what happens, if this usage is impaired. This can in the end not only help to better understand and fight frequent pathogens, but also to solve the complex puzzle of microbial interactions, niche formation, infection and resistance in the intestine, that is still at the beginning of being understood.

It is quite remarkable that the lack of pyruvate uptake has consequences not only for the utilization of this primary metabolite, but also for chemotaxis and survival in oxidative, nitrosative, and antibiotic stress in *S.* Typhimurium. On the other hand, compared with the wild-type, the pyruvate transporter deletion mutant had a more moderate disadvantage in survival in macrophages or in colonization of the mouse intestine and systemic infection. The *in vivo* results reflect the complexity of the gut ecosystem and the diversity of factors leading to colonization and infection by pathogens such as *S.* Typhimurium. However, it is the transport proteins in particular that play a very crucial role in microbial communities, allowing cross-feeding, but also achieving specificity as to which bacterium takes up which metabolite. This in turn contributes to community structure (Girinathan et al., 2021; Pontrelli et al., 2022; Weiss et al., 2021).

We and others have characterized pyruvate uptake systems in various gamma-proteobacteria. However, it remains unclear why different bacteria have different numbers of transporters and sensing systems. For *E. coli*, two pyruvate sensing systems and three pyruvate transporters (BtsT, YhjX, and CstA) were identified (Gasperotti et al., 2020), whereas for *S.* Typhimurium, only one pyruvate sensing system and two pyruvate transporters (BtsT and CstA) were found.

In contrast, the fish pathogen *Vibrio campbellii*, which excretes extraordinarily high amounts of pyruvate, harbors only one pyruvate sensing system and one transporter (Göing et al., 2021). Moreover, in contrast to pyruvate uptake systems, no exporter of pyruvate is known in any organism. As numerous bacteria excrete pyruvate, Tremblay et al. (2021) hypothesized that members of the gut microbiota might excrete pyruvate as a result of overflow metabolism, which then promotes the persistence of pathogens in the intestine. This metabolic cross-feeding of pyruvate was recently shown in another specific microbial community (Pontrelli et al., 2022). To gain more detailed knowledge of frequent pathogens on a molecular level regarding sensing systems, transporters, and their biological relevance might at some point tip the scales to understand the underlying functional structures and overcome worldwide burdens, such as severe gastroenteritis.

## EXPERIMENTAL PROCEDURES

### Strains, plasmids and oligonucleotides

*S.* Typhimurium and *E. coli* strains as well as plasmids used in this study are listed in Table 1. Oligonucleotide sequences are listed in Table S1. Molecular methods followed standard protocols (Sambrook et al., 1989) or were implemented according to manufacturer’s instructions.

**Table 1.**
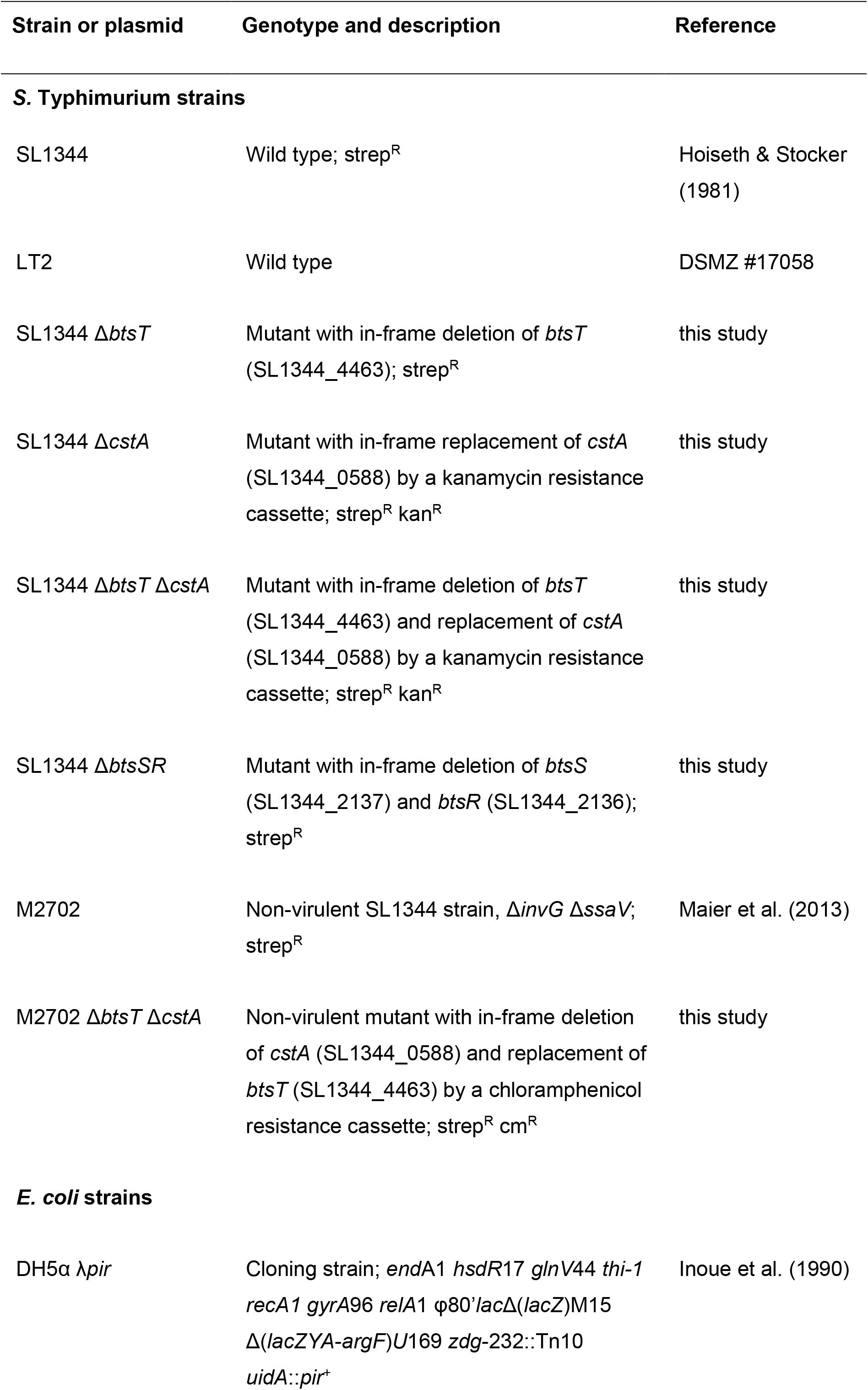
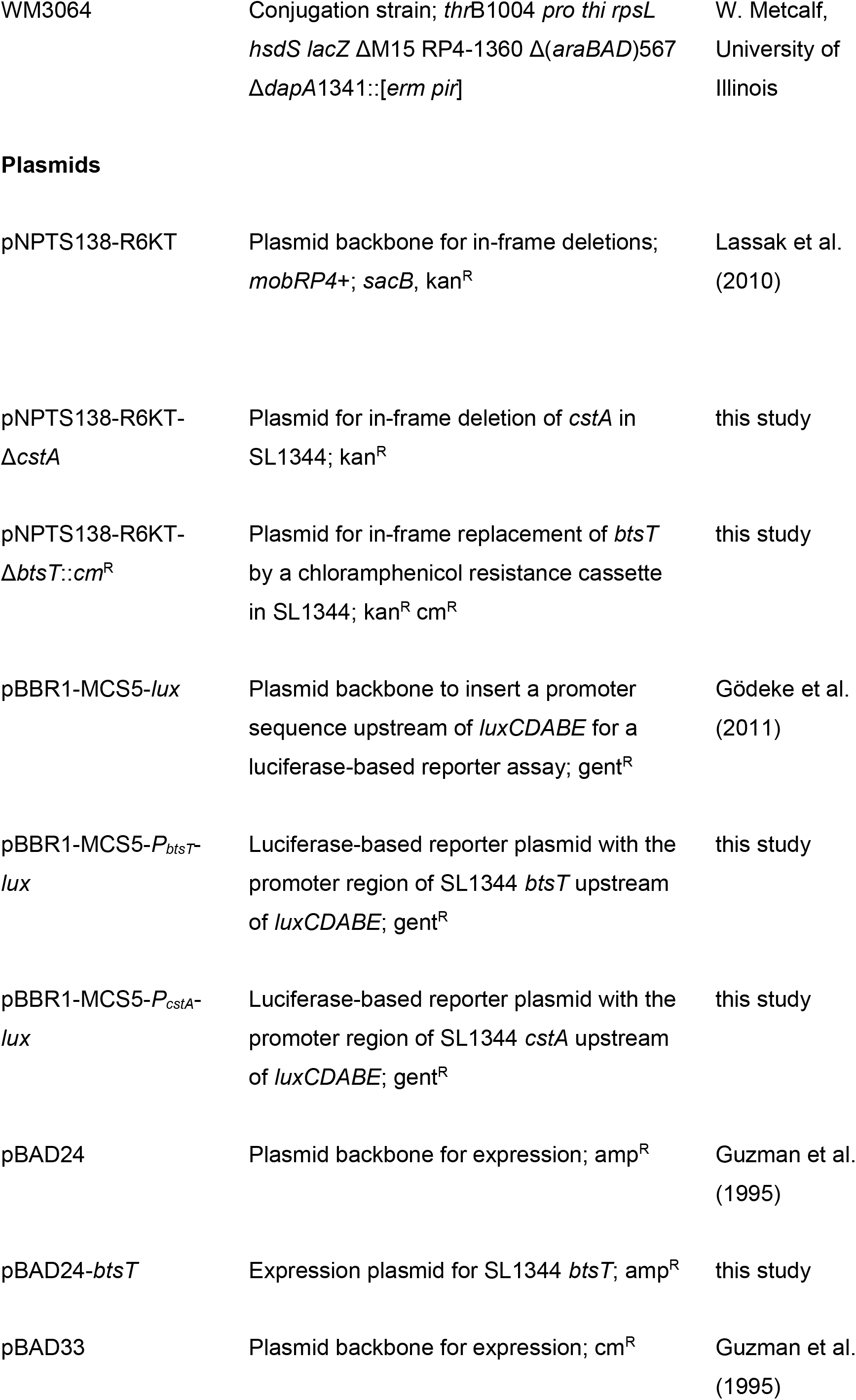
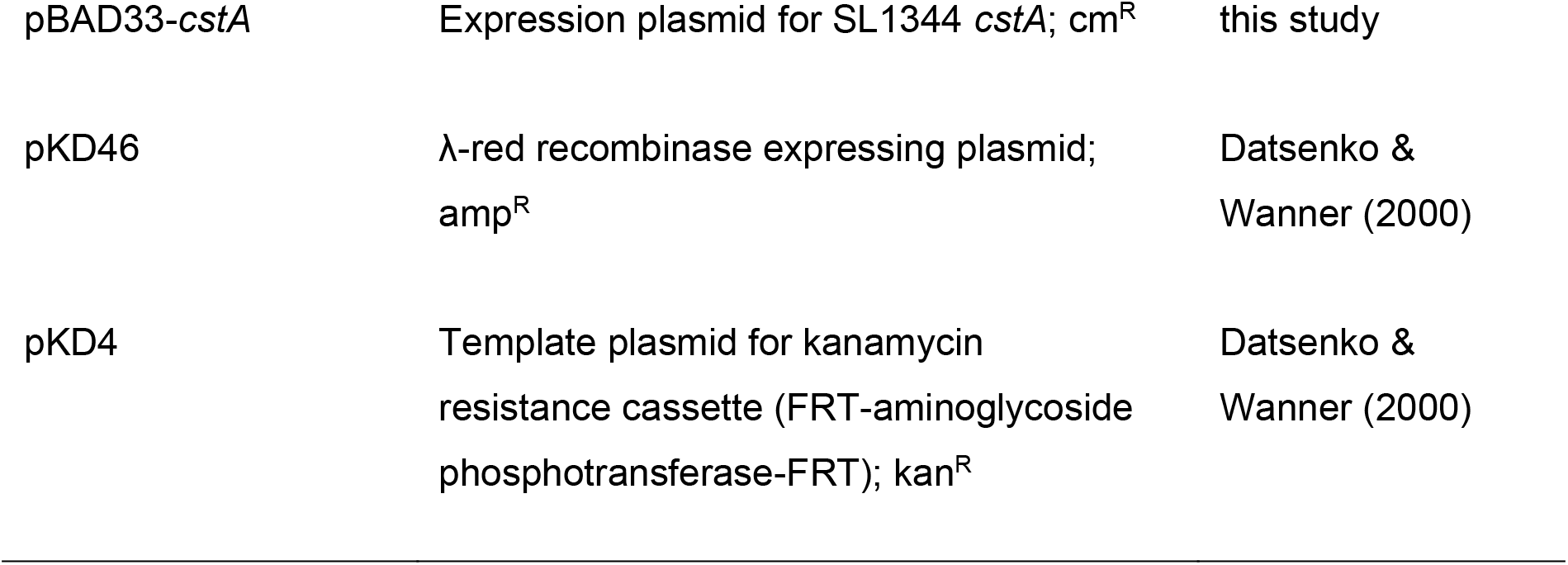
Strains and plasmids used in this study.

| Strain or plasmid | Genotype and description | Reference |
| --- | --- | --- |
| **S. Typhimurium strains** |  |  |
| SL1344 | Wild type; strep<sup>R</sup> | Hoiseth & Stocker (1981) |
| LT2 | Wild type | DSMZ #17058 |
| SL1344 $\Delta btsT$ | Mutant with in-frame deletion of *btsT* (SL1344_4463); strep<sup>R</sup> | this study |
| SL1344 $\Delta cstA$ | Mutant with in-frame replacement of *cstA* (SL1344_0588) by a kanamycin resistance cassette; strep<sup>R</sup> kan<sup>R</sup> | this study |
| SL1344 $\Delta btsT \Delta cstA$ | Mutant with in-frame deletion of *btsT* (SL1344_4463) and replacement of *cstA* (SL1344_0588) by a kanamycin resistance cassette; strep<sup>R</sup> kan<sup>R</sup> | this study |
| SL1344 $\Delta btsSR$ | Mutant with in-frame deletion of *btsS* (SL1344_2137) and *btsR* (SL1344_2136); strep<sup>R</sup> | this study |
| M2702 | Non-virulent SL1344 strain, $\Delta invG \Delta ssaV$; strep<sup>R</sup> | Maier et al. (2013) |
| M2702 $\Delta btsT \Delta cstA$ | Non-virulent mutant with in-frame deletion of *cstA* (SL1344_0588) and replacement of *btsT* (SL1344_4463) by a chloramphenicol resistance cassette; strep<sup>R</sup> cm<sup>R</sup> | this study |
| ***E. coli* strains** |  |  |
| DH5α $\lambda pir$ | Cloning strain; *endA1 hsdR17 glnV44 thi-1 recA1 gyrA96 relA1* $\phi 80' lac\Delta(lacZ)M15 \Delta(lacZYA-argF)U169 zdg-232::Tn10 uidA::pir^+$ | Inoue et al. (1990) |
| WM3064 | Conjugation strain; *thrB1004 pro thi rpsL hsdS lacZ* ΔM15 RP4-1360 $\Delta$(*araBAD*)567 Δ*dapA1341::[erm pir]* | W. Metcalf,<br>University of<br>Illinois |
| Plasmids |  |  |
| pNPTS138-R6KT | Plasmid backbone for in-frame deletions; *mobRP4+*; *sacB*, *kan*<sup>R</sup> | Lassak et al.<br>(2010) |
| pNPTS138-R6KT-Δ*cstA* | Plasmid for in-frame deletion of *cstA* in SL1344; *kan*<sup>R</sup> | this study |
| pNPTS138-R6KT-*ΔbtsT::cm*<sup>R</sup> | Plasmid for in-frame replacement of *btsT* by a chloramphenicol resistance cassette in SL1344; *kan*<sup>R</sup> *cm*<sup>R</sup> | this study |
| pBBR1-MCS5-*lux* | Plasmid backbone to insert a promoter sequence upstream of *luxCDABE* for a luciferase-based reporter assay; *gent*<sup>R</sup> | Gödeke et al.<br>(2011) |
| pBBR1-MCS5-*P*<sub>*btsT*</sub>-*lux* | Luciferase-based reporter plasmid with the promoter region of SL1344 *btsT* upstream of *luxCDABE*; *gent*<sup>R</sup> | this study |
| pBBR1-MCS5-*P*<sub>*cstA*</sub>-*lux* | Luciferase-based reporter plasmid with the promoter region of SL1344 *cstA* upstream of *luxCDABE*; *gent*<sup>R</sup> | this study |
| pBAD24 | Plasmid backbone for expression; *amp*<sup>R</sup> | Guzman et al.<br>(1995) |
| pBAD24-*btsT* | Expression plasmid for SL1344 *btsT*; *amp*<sup>R</sup> | this study |
| pBAD33 | Plasmid backbone for expression; *cm*<sup>R</sup> | Guzman et al.<br>(1995) |
| pBAD33-*cstA* | Expression plasmid for SL1344 *cstA*; $\text{cm}^R$ | this study |
| pKD46 | λ-red recombinase expressing plasmid;<br>$\text{amp}^R$ | Datsenko &<br>Wanner (2000) |
| pKD4 | Template plasmid for kanamycin<br>resistance cassette (FRT-aminoglycoside<br>phosphotransferase-FRT); $\text{kan}^R$ | Datsenko &<br>Wanner (2000) |

*S*. Typhimurium SL1344 mutants were first generated in strain LT2 and then transduced with phage P22 to strain SL1344. Clean in-frame deletions of *btsT* and *btsSR* in SL1344 were created by λ-Red recombination (Karlinsey, 2007). One-step inactivation of *cstA* by insertion of a chromosomal kanamycin resistance cassette with flanking regions (FRT-aminoglycoside phosphotransferase-FRT) was performed as described by Datsenko & Wanner (2000). Gene deletions were checked by colony PCR and confirmed by sequencing.

In *S*. Typhimurium M2702, clean in-frame deletion of *btsT* and gene inactivation of *cstA* by a chloramphenicol resistance cassette were performed by double homologous recombination using the pNPTS138-R6KT suicide plasmid as previously described (Brameyer et al., 2020; Lassak et al., 2010). *E. coli* DH5α *λpir* cells were used for cloning. Plasmid sequences were confirmed by sequencing and transferred into *S.* Typhimurium by conjugation using the *E. coli* WM3064 strain. Double homologous recombination was induced as described before (Göing et al., 2021). First, mutants with single-crossover integrations of the whole plasmid were selected on LB agar plates containing kanamycin. Then, the second crossover was induced by addition of 10% (wt/vol) sucrose and kanamycin-sensitive clones were checked by colony PCR. Gene deletions were confirmed by sequencing.

Complementation of deletion mutants was achieved by expressing the genes from plasmids. To this end, *btsT* and *cstA* were each amplified by PCR from SL1344 genomic DNA and cloned into plasmids pBAD24 and pBAD33, respectively, using restriction enzymes EcoRI and HindIII. Plasmids were transferred into the mutant strains by electroporation and leakiness of the arabinose promoter was sufficient for expression.

### Growth conditions

*S.* Typhimurium and *E. coli* strains were grown overnight under agitation (200 rpm) at 37°C in LB medium (10 g/l tryptone, 5 g/l yeast extract, 10 g/l NaCl). The conjugation strain *E. coli* WM3064 was grown in the presence of 300 μM diaminopimelic acid. If necessary, media were supplemented with 50 μg/ml kanamycin sulfate, 100 μg/ml ampicillin sodium salt, 30 μg/ml chloramphenicol and/or 20 μg/ml gentamicin sulfate to maintain plasmid(s) in the cells. To measure growth of *S.* Typhimurium strains on different carbon sources, cells were cultivated for 24 h at 37°C in M9 minimal medium (Harwood & Cutting, 1990) supplemented with 4 μg/ml histidine and the C-sources as indicated. Growth was monitored by measuring the optical density at 600 nm (OD_600_) over time.

### Luciferase reporter assay for the analysis of *btsT* and *cstA* expression

Expression of *btsT* and *cstA* was determined using a luciferase-based reporter assay. Reporter plasmids for *btsT* or *cstA* expression (pBBR1-MCS5-*PbtsT*-*lux* or pBBR1-MCS5-*PcstA*-*lux*) were constructed: Promoter regions of *btsT* and *cstA* (500 bp upstream of the start codon) were each amplified by PCR from SL1344 genomic DNA and cloned into the pBBR1-MCS5-*lux* vector, using restriction enzymes XbaI and XhoI. Plasmids were transferred into *S.* Typhimurium strains by electroporation. Cells harboring the reporter plasmid were grown in various media in 96-well-plates, inoculated from overnight cultures to a starting OD_600_ of 0.05. Plates were then incubated under constant agitation at 37°C and OD_600_ as well as luminescence values were measured at intervals of 10 min for 24 h in a ClarioStar plate reader (BMG). Gene expression was presented in relative light units (RLU) normalized to OD_600_.

### External pyruvate determination

Levels of excreted pyruvate were measured using a procedure adapted from O’Donnell-Tormey et al. (1987). *S.* Typhimurium strains were grown under agitation at 37°C in LB and growth was monitored. At selected time points, 1-ml samples of supernatant were harvested by centrifugation at 4°C (10 min, 14,000 x *g*). Proteins were precipitated by the addition of 250 μl ice-cold 2 M perchloric acid. After a 5-min incubation on ice, the samples were neutralized with 250 μl 2.5 M potassium bicarbonate, and precipitates were removed by centrifugation (4°C, 10 min, 14,000 x *g*). Pyruvate concentrations of the clear supernatants, diluted 1:5 in 100 mM PIPES buffer (pH 7.5), were determined using an enzymatic assay based on the conversion of pyruvate and NADH + H^+^ to lactate by lactate dehydrogenase. The assay was performed as described before (Gasperotti et al., 2020).

### Pyruvate uptake measurement

To determine the uptake of pyruvate by *S.* Typhimurium, a transport assay was performed with radiolabeled pyruvate. Cells were grown under agitation at 37°C in LB and harvested in mid-log phase. Cells were pelleted at 4°C, washed twice and resuspended in transport buffer (1 g/l (NH_4_)_2_SO_4_, 10 g/l K_2_HPO_4_, 4.5 g/l KH2PO4, 0.1 g/l MgSO4, pH 6.8) to an absorbance of 5 at 420 nm, equivalent to a total protein concentration of 0.35 mg/ml. Uptake of ^14^C-pyruvate (55 mCi/mmol, Biotrend) was measured at a total substrate concentration of 10 μM at 18°C. At various time intervals, transport was terminated by the addition of ice-cold stop buffer (100 mM potassium phosphate, pH 6.0, 100 mM LiCl) followed by rapid filtration through membrane filters (MN gf-5, 0.4 μm nitrocellulose, Macherey Nagel). The filters were dissolved in 5 ml scintillation fluid (MP Biomedicals), and radioactivity was determined in a liquid scintillation analyzer (Perkin-Elmer).

### Motility assay

Overnight cultures of *S.* Typhimurium were adjusted to an OD_600_ of 1 and 10 μl were inoculated into freshly poured swimming motility plates (10 g/l tryptone, 5 g/l NaCl, 0.3% agar, wt/vol) and incubated at 37°C for 3 h. Pictures were taken with a Canon EOS M50 camera and images were analyzed using the software ImageJ (Schneider et al., 2012). The size of the ring was measured and the size of each ring was expressed relatively to the average size of the wild-type ring.

### Chemotaxis test

Chemotaxis of *S.* Typhimurium towards different compounds was tested using the plug-in-pond assay (Darias et al., 2014). Cells grown in LB were pelleted, resuspended to a final OD_600_ of 0.4 in M9 soft agar (M9 medium with 0.3% agar wt/vol), and poured into a petri dish, in which agar plugs (M9 medium with 1.5% agar, w/v) containing the test substances had been placed. Plates were incubated at 37°C for 3 h. Pictures were taken with a Canon EOS M50 camera.

### Stress assay

To test survival under oxidative and nitrosative stress, *S.* Typhimurium cells were grown in LB to an OD_600_ of 1.2, split into groups and either treated with 12.5 mM H_2_O_2_ for H_2_O_2_ stress, 250 μM spermine NONOate for NO stress or with H_2_O as a control. After 20 min incubation, catalase was added (for H_2_O_2_ only), and cells were plated in dilutions on LB to determine CFU. Survival under stress was calculated as the percentage of CFU in relation to the control condition and wild type values were set to 100%.

### Persister formation

To investigate persister formation, *S.* Typhimurium cells were grown in LB to an OD_600_ of 1.2 and diluted to an OD_600_ of 0.05 into fresh LB containing 50 μg/ml gentamicin. Every hour, cells were plated in dilutions on LB agar plates to determine CFU, which represent cells being able to survive the antibiotic treatment by forming persister cells.

### Intramacrophage antibiotic survival assays

*S.* Typhimurium strains were grown in LB for 16 h. Stationary phase bacteria were opsonized with 8% (wt/vol) mouse serum (Sigma) for 20 min and added to the bone marrow derived macrophages at a multiplicity of infection (MOI) of 5. Infection was then synchronized by 5 min centrifugation at 100 x *g.* The infected macrophages were incubated for 30 min at 37°C with 5% CO_2_ to allow phagocytosis to occur. At 30 min following infection, the macrophages were washed three times with PBS and half of the cells were lysed with 0.1% (vol/vol) Triton X-100 in PBS. Bacteria were then centrifuged at 16,000 x *g* for 2 min at room temperature, following resuspension in PBS. The bacteria were diluted ten-fold in PBS and plated on LB agar to count the number of CFU prior to antibiotic treatment. With regards to the remaining macrophages, the three PBS washes were followed by addition of fresh medium (Dulbecco’s modified eagle medium with high glucose (DMEM), 10% (vol/vol) fetal calf serum, 10 mM HEPES, 1 mM sodium pyruvate) containing 100 μg/ml cefotaxime. Cefotaxime was added to test intramacrophage antibiotic survival for 24 h. At 24 h following antibiotic treatment, the cells were washed three times with PBS, then lysed with 0.1% (vol/vol) Triton X-100 in PBS. Bacteria were then centrifuged at 16,000 x *g* for 2 min at room temperature, following resuspension in PBS. The bacteria were diluted ten-fold in PBS and plated on LB agar to count the number of CFU following antibiotic treatment. 24 h survival was expressed as a fold change of wild-type values.

### Infection of gnotobiotic mice

All animal experiments were approved by the local authorities (Regierung von Oberbayern). Germ-free C57BL/6J mice and C57BL/6J mice colonized with defined bacterial consortia (OMM^12^) were obtained from the animal housing facility of the Max von Pettenkofer-Institute (Ludwig-Maximilians-University, Munich, Germany). Mice were housed under germfree conditions in flexible film isolators (North Kent Plastic Cages) or in Han-gnotocages (ZOONLAB). The mice were supplied with autoclaved ddH_2_O and Mouse-Breeding complete feed for mice (Ssniff) ad libitum. For all experiments, female and male mice between 6-15 weeks were used, and animals were randomly assigned to experimental groups. Mice were not single-housed and kept in groups of 3-5 mice per cage during the experiment. All animals were scored twice daily for their health status.

For generation of the ASF mouse line, germfree C57BL/6J mice were inoculated with a mixture of ASF^3^ (ASF356, ASF361, ASF519). Mice were inoculated twice (72 h apart) with the bacterial mixtures (frozen glycerol stocks) by gavage (50 μl orally, 100 μl rectally). Mice were housed under germfree conditions and were used 12 days post inoculation for experiments to ensure stable colonization of the consortium.

For infection experiments with virulent *S.* Typhimurium SL1344, OMM^12^ mice were treated with streptomycin by oral gavage with 50 μl of 500 mg/ml streptomycin one day before infection. For infection experiments with avirulent *S.* Typhimurium M2702, OMM^12^ and ASF^3^ mice were not treated with streptomycin before infection. For all infection experiments, both *S.* Typhimurium wild type and mutant cells were grown on MacConkey agar plates (Oxoid) containing streptomycin (50 mg/ml) at 37°C. One colony was re-suspended in 5 ml LB containing 0.3 M NaCl and grown for 12 h at 37°C on a wheel rotor. A subculture (1:20 dilution) was prepared in fresh LB containing 0.3 M NaCl and incubated for further 4h. Bacteria were washed with ice-cold sterile PBS, pelleted and re-suspended in fresh PBS. *S.* Typhimurium wild type and mutant cells were mixed in a 1:1 ratio adjusted by OD_600_. Mice were infected with the *S.* Typhimurium mix by oral gavage with 50 μl of bacterial suspension (approximately 4 x 10^6^ CFU).

*S.* Typhimurium total loads in feces were determined on the first day after infection by plating on MacConkey agar with streptomycin (50 mg/ml). All mice were sacrificed by cervical dislocation four days after infection, and *S.* Typhimurium total loads in fecal and cecal contents, as well as from lymph nodes, spleen and liver were determined by plating on MacConkey agar with streptomycin (50 mg/ml). From each plate 50 colonies were picked onto MacConkey agar plates with streptomycin (50 mg/ml) and chloramphenicol (30 mg/ml) for M2702 mutants or kanamycin (30 mg/ml) for SL1344 mutants to determine the competitive index between wild type and mutant strain.

## ACKNOWLEDGEMENTS

The authors thank Raphaela Götz for her contribution in strain construction. This research was funded by the Deutsche Forschungsgemeinschaft (project number 395357507-SFB1371 to K.J. and B.S.).

## AUTHOR CONTRIBUTIONS

Authors made substantial contributions in the following aspects: SP: conceptualization, data acquisition/analysis/interpretation and writing; FF, ASW and AM: data acquisition/analysis; SH and BS: conceptualization, data analysis/interpretation; KJ: conceptualization, data analysis/interpretation and writing.

