## Supplementary material for "Identification of two pyruvate transporters in *Salmonella enterica* serovar Typhimurium and their biological relevance": Table S1

[2] Present affiliation: Bayer AG, Berlin, Germany

### SUPPLEMENTAL METHODS

**Strain construction of SL1344 *btsT*::*mNeonGreen*.** Chromosomal fusions were created by double homologous recombination using the pNPTS138-R6KT suicide plasmid (Lassak et al., 2010). SL1344 *btsT* was amplified by PCR from SL1344 genomic DNA with oligonucleotides #38 and #39, without keeping the stop codon. The *mNeonGreen* gene was amplified by PCR from a plasmid (Peter Graumann, Marburg) with oligonucleotides #40 and #41. An 800 bp region directly downstream of *btsT* was amplified by PCR from SL1344 genomic DNA with oligonucleotides #42 and #43. The pNPTS138-R6KT backbone was linearized with oligonucleotides #37 and #44. All fragments were created with overlaps of 20 bp for assembly using the NEBuilder kit (New England Biolabs). The final pNPTS138-R6KT-*btsT*::*mNeonGreen* plasmid was first transformed into *E. coli* DH5α and confirmed by sequencing. Then it was transferred into *E. coli* WM3064 for conjugation with SL1344. Double homologous recombination was induced as described in the main manuscript. Oligonucleotide sequences are listed in table S1.

***In vivo* single cell fluorescence measurements.** SL1344 cells with the chromosomal fusion *btsT*::*mNeonGreen* were grown in InSPI2 and NonSPI2 medium (Löber et al., 2006) with 4 µg/ml histidine and 60 mM pyruvate as carbon source, inoculated from overnight culture to an initial $OD_{600}$ of 0.05. In exponential growth phase, samples were taken and 2 µl of the culture were spotted on 1% (wt/vol) agarose prepared with PBS on a microscope slide and sealed with a cover slide. Microscopy was performed using a Leica DMI6000 B fluorescence microscope, with an excitation wavelength of 485 nm and a 510-nm emission filter. To quantify single cell fluorescence, 1000 cells per condition (InSPI2 or NonSPI2) were analyzed using the plug-in MicrobeJ (Ducret et al., 2016) of the software ImageJ (Schneider et al., 2012), as described before (Brameyer et al., 2020). The background was subtracted for each cell.

**SUPPLEMENTAL FIGURES**

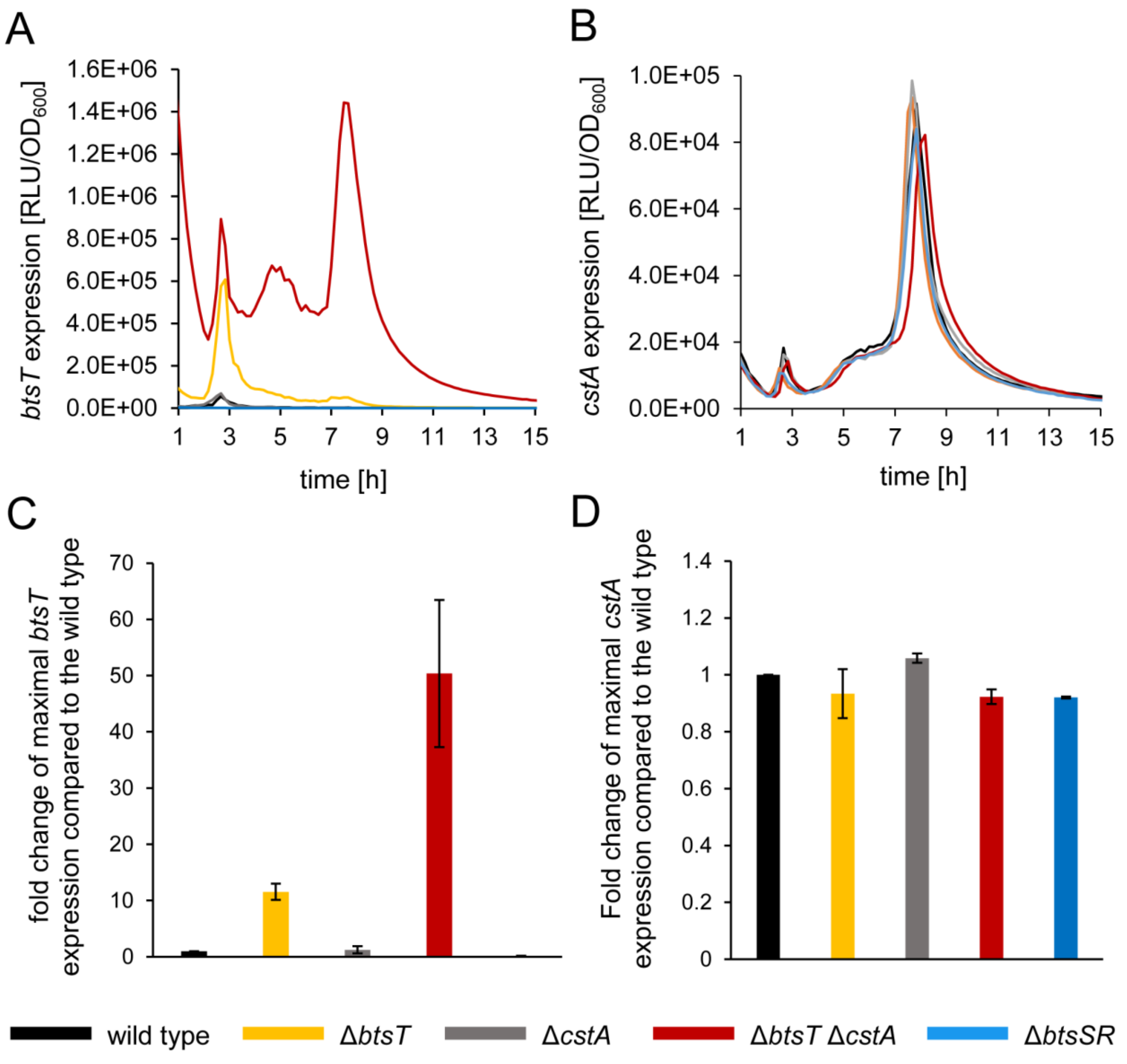


**Figure S1. Expression of *btsT* and *cstA* in *S*. Typhimurium mutants.** SL1344 wild-type (black), Δ*btsT* (yellow), Δ*cstA* (grey), Δ*btsT*Δ*cstA* (red) and Δ*btsSR* (blue) cells harboring the reporter plasmid for *btsT* (pBBR1-MCS5-$P_{btsT}$-*lux*) or for *cstA* (pBBR1-MCS5-$P_{cstA}$-*lux*) were grown in LB medium in a plate reader at 37°C. Luminescence values were measured over time and gene expression was determined as RLU per $OD_{600}$. **A)** Expression of *btsT* during growth. **B)** Expression of *cstA* during growth. **C)** Maximal *btsT* expression depicted as fold change from wild-type value. **D)** Maximal *cstA* expression depicted as fold change from the wild-type value. A, B: graphs represent the mean of three independent replicates. the standard deviations were below 10%. C, D: Error bars represent the standard deviations of the mean of three independent replicates.

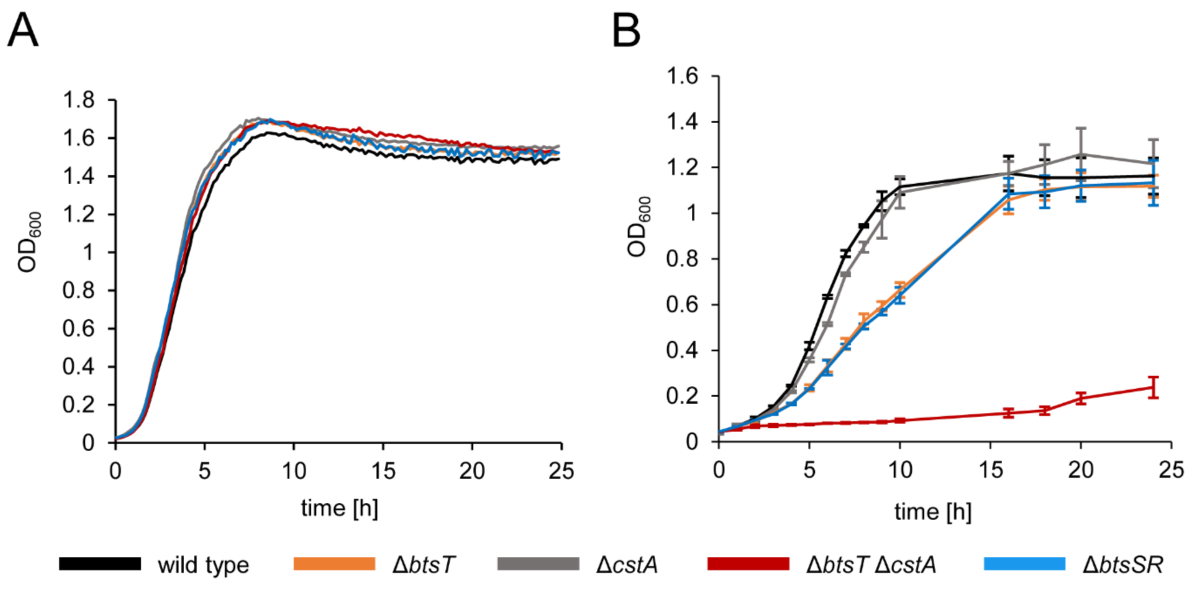


**Figure S2. Growth of *S.* Typhimurium mutants.** SL1344 wild-type (black), Δ*btsT* (yellow), Δ*cstA* (grey), Δ*btsT*Δ*cstA* (red) and Δ*btsSR* (blue) cells were grown for 24 hours at 37°C in **A)** LB medium or **B)** M9 minimal medium with 60 mM pyruvate.

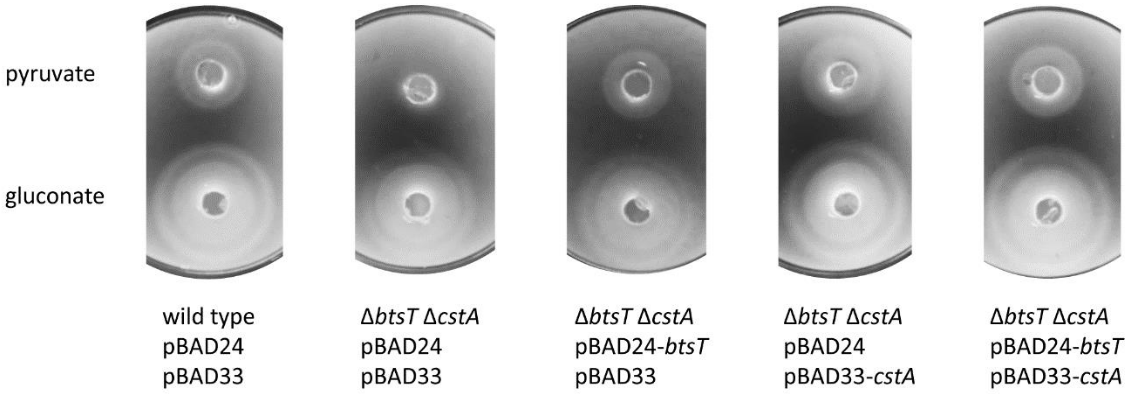


**Figure S3. *S*. Typhimurium Δ*btsT*Δ*cstA* mutant lost chemotactic response to pyruvate.** Chemotaxis was tested by mixing SL1344 wild-type or Δ*btsT*Δ*cstA* cells harboring the indicated expression plasmids for *btsT* (pBAD24-*btsT*) and/or *cstA* (pBAD33-*cstA*) or the empty vectors with 0.3% (wt/vol) M9 soft agar and pouring them over 1.5% (wt/vol) M9 agar plugs containing either 60 mM gluconate (above) or 60 mM pyruvate (below). Plates were incubated at 37°C for 4 hours, and the pictures are representative of three independent experiments.

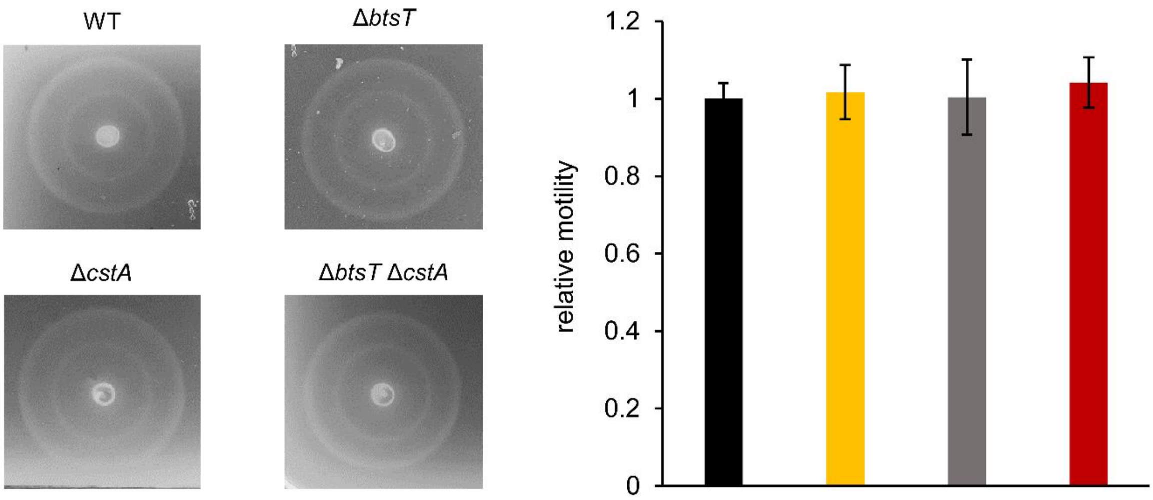


**Figure S4. Motility of *S*. Typhimurium is not affected by deletions of *btsT* or *cstA*.** Motility of SL1344 wild type (upper left, black), Δ*btsT* (upper right, yellow), Δ*cstA* (lower left, grey) and Δ*btsT*Δ*cstA* (lower right, red) cells was tested by spotting equal numbers of cells on 0.3% LB soft agar, incubating the plates at 37°C for 3 hours and measuring the diameter of the ring with the software ImageJ. Images of rings are representative of four independent experiments and relative motility was determined in relation to the mean diameter of the wild type ring.

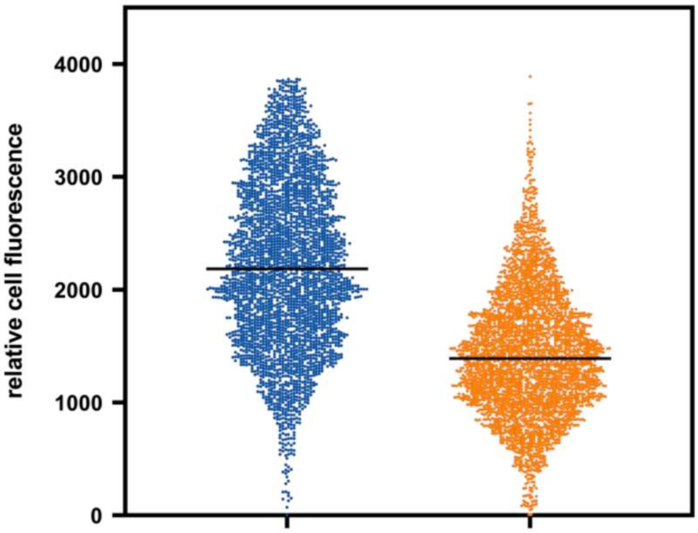


**Figure S5. Expression of *btsT* in *S*. Typhimurium under SPI2 inducing conditions.** Strain SL1344 *btsT*::*mNeonGreen*, which chromosomally encodes a fusion between *btsT* and *mNeonGreen,* was used to measure expression of *btsT* in single cells. Cells were grown in NonSPI2 (blue) or InSPI2 (orange) medium with 4 µg/ml histidine and 60 mM pyruvate as C-source until mid-exponential phase. Samples were taken for fluorescence microscopy. To quantify relative fluorescence intensities of single cells, phase contrast and fluorescent images were analyzed using the ImageJ (Schneider et al., 2012) plugin MicrobeJ (Ducret et al., 2016). In total >1000 cells were quantified per condition.

**SUPPLEMENTAL TABLES**

**TABLE S1.** Oligonucleotides used in this study

| | **DNA sequence** | **Description** |
|---|---|---|
| #1 | CGCTGTGCGCTACCCGGCATCAGTTTTAGTGGTGTGAAGATTAAGACCCACTTTCACATT | Primer 1 for tetRA-insertion by λ-Red recombination in *btsT* |
| #2 | ATTAAACTTACAACCAGGTTTTACTATGGATACGAAAAAGCTAAGCACTTGTCTCCTG | Primer 2 for tetRA-insertion by λ-Red recombination in *btsT* |
| #3 | ATTAAACTTACAACCAGGTTTTACTATGGATACGAAAAAGTCTTCACACCACTAAAACTG | Primer 1 for clean deletion of *btsT* |
| #4 | AGTCCGGAATACCAATCAACA | Primer 2 for clean deletion of *btsT* |
| #5 | ACCGCTTAAACCGCCATACA | Primer 1 for sequencing of Δ*btsT* |
| #6 | ACGTTCGCGGAAGAACTCTT | Primer 2 for sequencing of Δ*btsT* |
| #7 | GGCAAAACGATATTCTAACAGTCTTTTACAGGCCAATCGCTTAAGACCCACTTTCACATT | Primer 1 for tetRA-insertion by λ-Red recombination in *btsSR* |
| #8 | TTTAATTGAAGTGTGGTTTGCGGGTATGTACGAGTTTAATCTAAGCACTTGTCTCCTG | Primer 2 for tetRA-insertion by λ-Red recombination in *btsSR* |
| #9 | TTGTTGATACGACGTTCCGC | Primer 1 for clean deletion of *btsSR* |
| #10 | TTTAATTGAAGTGTGGTTTGCGGGTATGTACGAGTTTAATGCGATTGGCCTGTAAAAGAC | Primer 2 for clean deletion of *btsSR* |
| #11 | TGGAACACCCAAACGGACAACAACTATGAATAAATCAGGGTAGGCTGGAGCTGCTTCGAA | Primer 1 to amplify the FRT-kanamycin-FRT cassette from pKD46 for replacement of *cstA* |
| #12 | GGAGAGGGCTATTGATGTAAAAAGATTAGTGCGCGCCTTTTCCTCCTTAGTTCCTATTCC | Primer 2 to amplify the FRT-kanamycin-FRT cassette from pKD46 for replacement of *cstA* |
| #13 | CTCTTTGACGAGCAGGGGAG | Primer 1 for sequencing of Δ*cstA* |
| #14 | CGTCTGATCCGGATGCGTTA | Primer 2 for sequencing of Δ*cstA* |
| #15 | AAAAAATCTAGAGCGATGACGTGCTGGAGGCG | Primer 1 to amplify the promoter of *cstA* to create pBBR1-MCS5-$P_{cstA}$-*lux*, XbaI site |
| #16 | AAAAAACTCGAGAGTTGTTGTCCGTTTGGGTG | Primer 2 to amplify the promoter of *cstA* to create pBBR1-MCS5-$P_{cstA}$-*lux*, XhoI site |
| #17 | AAAAAATCTAGAAGTTTGCAATACGGTGAAGT | Primer 1 to amplify the promoter of *btsT* to create pBBR1-MCS5-$P_{btsT}$-*lux*, XbaI site |
| #18 | AAAAAACTCGAGAGTAAAACCTGGTTGTAAGT | Primer 2 to amplify the promoter of *btsT* to create pBBR1-MCS5-$P_{btsT}$-*lux*, XhoI site |
| #19 | GCGCGCGAATTCACTATGGATACGAAAAAGATATT | Primer 1 to amplify *btsT* to create pBAD24-*btsT*, EcoRI site |
| #20 | GCTAGCAAGCTTTTAGTGATGGTGATGGTGATGGTGGTGTGAAGAGATCTTCA | Primer 2 to amplify *btsT* to create pBAD24-*btsT*, HindIII site |
| #21 | GCGCGCGAATTCACTATGAATAAATCAGGGAAATA | Primer 1 to amplify *cstA* to create pBAD3-*cstA*, EcoRI site |
| #22 | GCTAGCAAGCTTTTAGTGATGGTGATGGTGATGGTGCGCGCCTTTCGCCTGCG | Primer 2 to amplify *cstA* to create pBAD3-*cstA*, HindIII site |

| | | |
|---|---|---|
| #23 | ACTCACCCATAACGTGCGCTGCATACAG ATATCCTGCAGAGAAGCTTGGC | Primer 1 to amplify pNPTS138-R6KT to create pNPTS138-R6K-Δ*cstA* for clean deletion of *cstA* |
| #24 | GCCAAGCTTCTCTGCAGGATATCTGTAT GCAGCGCACGTTATGGGTGAGT | Primer 1 to amplify 800 bp upstream region of *cstA* to create pNPTS138-R6K-Δ*cstA* for clean deletion of *cstA* |
| #25 | GGAGAGGGCTATTGATGTAAAAAGAAGT TGTTGTCCGTTTGGGTGTTCCA | Primer 2 to amplify 800 bp upstream region of *cstA* to create pNPTS138-R6K-Δ*cstA* for clean deletion of *cstA* |
| #26 | TGGAACACCCAAACGGACAACAACTTCT TTTTACATCAATAGCCCTCTCC | Primer 1 to amplify 800 bp downstream region of *cstA* to create pNPTS138-R6K-Δ*cstA* for clean deletion of *cstA* |
| #27 | GCCGAAGCTAGCGAATTCGTGGATCGTA AGGCTGGGCATTAACAGCGCGT | Primer 2 to amplify 800 bp downstream region of *cstA* to create pNPTS138-R6K-Δ*cstA* for clean deletion of *cstA* |
| #28 | ACGCGCTGTTAATGCCCAGCCTTACGAT CCACGAATTCGCTAGCTTCGGC | Primer 1 to amplify pNPTS138-R6KT to create pNPTS138-R6K-Δ*cstA* for clean deletion of *cstA* |
| #29 | TTATTCAGCTTCTTGGCGCGGGTAACAG ATATCCTGCAGAGAAGCTTGGC | Primer 1 to amplify pNPTS138-R6KT to create pNPTS1138-R6K-Δ*btsT*::cm$^R$ for replacement of *btsT* |
| #30 | GCCAAGCTTCTCTGCAGGATATCTGTTAC CCGCGCCAAGAAGCTGAATAA | Primer 1 to amplify 800 bp upstream region of *btsT* to create pNPTS1138-R6K-Δ*btsT*::cm$^R$ for replacement of *btsT* |
| #31 | ACGGCAAAAGCACCGCCGGACATCAAGT AAAACCTGGTTGTAAGTTTAAT | Primer 2 to amplify 800 bp upstream region of *btsT* to create pNPTS1138-R6K-Δ*btsT*::cm$^R$ for replacement of *btsT* |
| #32 | CGATGAGTGGCAGGGCGGGGCGTAAAA CTGATGCCGGGTAGCGCACAGCG | Primer 1 to amplify 800 bp downstream region of *btsT* to create pNPTS1138-R6K-Δ*btsT*::cm$^R$ for replacement of *btsT* |
| #33 | CCGAAGCTAGCGAATTCGTGGATCTCGC CTGCCACGTCGGTTTTGGTCAA | Primer 2 to amplify 800 bp downstream region of *btsT* to create pNPTS1138-R6K-Δ*btsT*::cm$^R$ for replacement of *btsT* |
| #34 | ATTAAACTTACAACCAGGTTTTACTTGAT GTCCGGCGGTGCTTTTGCCGT | Primer 1 to amplify chloramphenicol resistance cassette from pBAD33 to create pNPTS1138-R6K-Δ*btsT*::cm$^R$ for replacement of *btsT* |
| #35 | CGCTGTGCGCTACCCGGCATCAGTTTTA CGCCCCGCCCTGCCACTCATCG | Primer 2 to amplify chloramphenicol resistance cassette from pBAD33 to create pNPTS1138-R6K-Δ*btsT*::cm$^R$ for replacement of *btsT* |
| #36 | TTGACCAAAACCGACGTGGCAGGCGAGA TCCACGAATTCGCTAGCTTCGG | Primer 2 to amplify pNPTS138-R6KT to create pNPTS1138-R6K-Δ*btsT*::cm$^R$ for replacement of *btsT* |
| #37 | GATACCTACTGCCAGCGTTGCAGATATC CTGCAGAGAAGCTTGGC | Primer 1 to amplify pNPTS138-R6KT to create pNPTS138-R6KT-*btsT*::*mNeonGreen* |
| #38 | GCTTCTCTGCAGGATATCTGCAACGCTG GCAGTAGGTATCGCGCA | Primer 1 to amplify *btsT* to create pNPTS138-R6KT-*btsT*::*mNeonGreen* |

| | | |
|---|---|---|
| #39 | ACCATAGAACCGCCGCCACCGTGGTGTG AAGAGATCTTCACGCCG | Primer 2 to amplify *btsT* without stop codon to create pNPTS138-R6KT-*btsT*::*mNeonGreen* |
| #40 | TGAAGATCTCTTCACACCACGGTGGCGG CGGTTCTATGGTGAGCA | Primer 1 to amplify mNeonGreen to create pNPTS138-R6KT-*btsT*::*mNeonGreen* |
| #41 | TGCGCTACCCGGCATCAGTTTTACTTGTA CAGCTCGTCCATGCCC | Primer 2 to to amplify mNeonGreen to create pNPTS138-R6KT-*btsT*::*mNeonGreen* |
| #42 | TGGACGAGCTGTACAAGTAAAACTGATG CCGGGTAGCGCACAGCG | Primer 1 to amplify 800 bp downstream flank of *btsT* to create pNPTS138-R6KT-*btsT*::*mNeonGreen* |
| #43 | AGCTAGCGAATTCGTGGATCGGGCTATG GTGAACTGATTCATCTG | Primer 2 to amplify 800 bp downstream flank of *btsT* to create pNPTS138-R6KT-*btsT*::*mNeonGreen* |
| #44 | GAATCAGTTCACCATAGCCCGATCCACG AATTCGCTAGCTTCGGC | Primer 2 to amplify pNPTS138-R6KT to create pNPTS138-R6KT-*btsT*::*mNeonGreen* |
